# Heterogeneous short-term plasticity reformats latent temporal information into an explicit population clock

**DOI:** 10.64898/2026.11.17.767281

**Authors:** Nathan P. Buerkle, James M. Jeanne

## Abstract

Short-term plasticity (STP) supports diverse synaptic-level computations, but how its organization across a circuit shapes emergent, network-level dynamics remains poorly understood. Here, we first characterize STP across sparse inputs to Kenyon cells in the *Drosophila* mushroom body, revealing an unstructured and near-random organization that could not be explained by either pre- or postsynaptic cell identity. Rather than representing biological noise, an experimentally constrained network model instead shows how this heterogeneous organization reformats temporal information embedded in presynaptic spike rate trajectories into a behaviorally relevant population clock. Although an ideal observer could recover the same degree of temporal information from most alternative models, only the STP-mediated transformation made that information accessible to a biologically realistic decoder. Together, our results identify heterogeneous STP as a distributed biophysical mechanism that collectively functions as a temporal basis set to support circuit computations across variable input statistics.

## INTRODUCTION

Neurons and synapses exhibit considerable heterogeneity in their physiological properties, and this diversity is increasingly recognized as a computational feature rather than biological noise^1,2^. Heterogeneity spans multiple levels of circuit organization, with neurons of the same cell type often differing in gene expression^3,4^, intrinsic properties^5,6^, synaptic dynamics^7–9^, and patterns of connectivity^10–12^. Theoretical and empirical work has linked these heterogeneities to a wide range of computational functions such as enhanced coding capacity, temporal processing, and improved learning^5,13–19^, while loss of heterogeneity has been associated with neurological disorders^20,21^. Different forms of heterogeneity may therefore contribute to neural computation in distinct ways, depending on the properties that vary and the circuits in which they are expressed.

Short-term plasticity (STP) is a ubiquitous feature of synapses that dynamically modulates synaptic efficacy on timescales of hundreds of milliseconds to seconds. Depressing and facilitating synapses represent two broadly defined phenotypes that serve distinct computational roles as low-pass and high-pass filters, respectively^22–24^. Heterogeneity has been observed in both cortical and cerebellar circuits, where STP phenotypes can show consistent, computationally relevant biases based on pre- and postsynaptic cell identity^8,9,25–28^. Similar to other forms of neuronal heterogeneity, however, STP also has substantial within-class variability, and the effects of this diversity on emergent network activity remain largely unexplored.

The *Drosophila* mushroom body functions as the primary site of olfactory associative learning and presents an anatomically tractable circuit for relating synaptic organization to population dynamics^29^. Odor stimuli are first encoded by 51 types of antennal lobe projection neuron (PN), which project to a much larger population of ∼2,000 mushroom body intrinsic Kenyon cells (KCs)^30^. PN-KC connectivity is sparse and divergent, with each KC pseudo-randomly connected to only 6 PNs on average^31,32^. Together with inhibitory input from the local neuron APL, this architecture forms a canonical pattern separation circuit where KCs encode odor identity through a sparse population code^33,34^. KC output then converges on mushroom body output neurons (MBONs), with synapses that undergo dopamine-mediated long-term plasticity to drive learned behavior^35,36^. The computations performed by the KC population therefore define the representations available to downstream MBONs.

Despite the relatively simple and largely feedforward anatomy, KC odor responses exhibit complex temporal dynamics that form a behaviorally relevant population clock^37^. KCs preferentially spike at different times to collectively tile temporal space, and this clock extends to stimuli at least 10 s in duration. PNs have stereotyped odor responses with no explicit representation of time that could directly explain these dynamics^38,39^. Because PNs provide the only source of fast excitation to KCs, this raises the question of what circuit mechanisms transform temporally unstructured PN activity into a structured KC population clock. The emergence of this temporal structure likely reflects coordinated interactions between synaptic dynamics, intrinsic KC properties, and inhibition.

Here we focused on the contribution of PN-KC synaptic dynamics to KC odor responses. We first experimentally characterized STP phenotypes across PNs converging onto the same KC, which revealed substantial heterogeneity that could not be explained by either PN or KC identity. These data were fit with the Tsodyks-Markram model of STP^40,41^, and parameter estimates were used to construct an experimentally constrained model of the mushroom body to determine how STP influences circuit computations. Only models with a heterogeneous organization of STP could reproduce KC odor responses observed *in vivo*, with STP functioning to transform latent temporal information embedded in PN spike rates into a structured temporal representation necessary for downstream decoding by an MBON. Thus, our results extend the functional role of STP from a local filter on synaptic transmission to a distributed computational resource through its organization across a circuit.

## RESULTS

### 2-photon optogenetic measurement of PN-KC synaptic dynamics

We measured STP at PN-KC synapses in an *ex vivo* preparation using KC electrophysiology combined with 2-photon optogenetic stimulation of PNs^42^ (Figures 1A-1C and S1). Experiments were performed in *Drosophila* with the genotype *GH146-Gal4* > *UAS-CsChrimson::mVenus*, providing both optogenetic control and fluorescent labeling in 37 of 51 glomeruli^43^. We recorded from 89 KCs, of which 27 KCs comprising 108 synapses across 33 glomeruli met our inclusion criteria and were used in the final dataset (STAR Methods).

**Figure 1.**
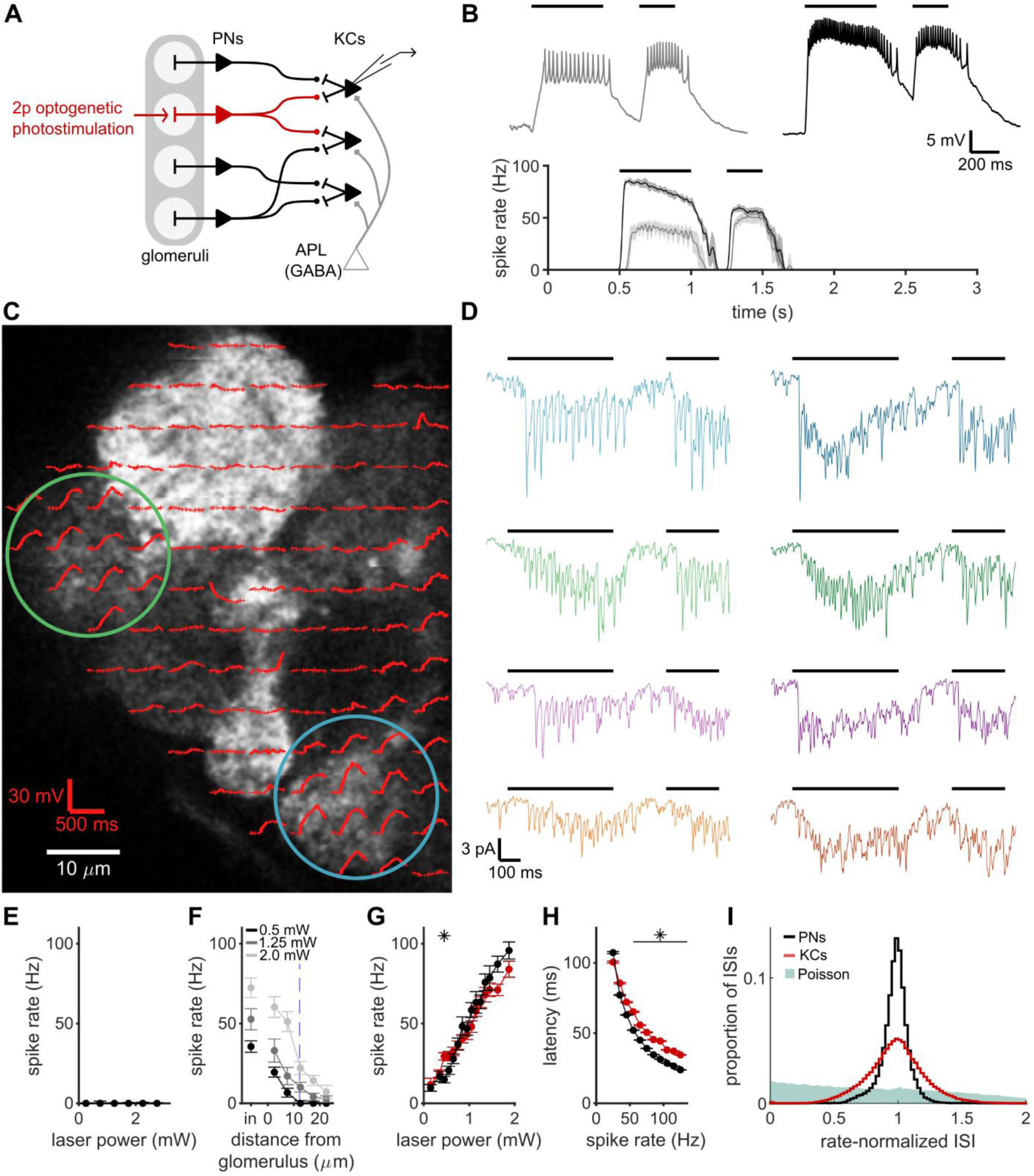
Measuring STP using 2-photon optogenetic stimulation of PN-KC synapses. **(A)** Schematic of mushroom body circuit and experimental design. **(B)** Example responses from a PN responding to 2-photon optogenetic stimulation at a low (left, 0.7 mW) and high (right, 1.8 mW) laser power. The bottom shows PSTHs averaged across trials with similar onset spike rates (mean ± SD, n = 10 trials). Black bars denote time of glomerular photostimulation. **(C)** Antennal lobe image via 2-photon microscopy overlaid with the voltage response of a γ KC to 2-photon optogenetic stimulation of spatially restricted voxels. Circles highlight two glomeruli that evoked consistent KC depolarizations. Connectivity mapping used a single 250 ms stimulus. **(D)** Example KC voltage-clamp recordings during 2-photon stimulation of 4 connected glomeruli. Left and right columns correspond to low and high laser powers, respectively. Rows 1 and 2 correspond to circled glomeruli in panel C. Glomeruli for rows 3 and 4 were located in different focal planes. **(E)** Control recordings from PNs (n = 15, mean ± SEM) demonstrate high lateral spatial resolution of 2-photon stimulation. Stimulation locations were selected 1-2 μm outside the target glomerulus, and spike rate was quantified at different laser powers. **(F)** Control recordings from PNs (n = 19, mean ± SEM) show axial spatial resolution of 2-photon stimulation. Stimulation sites were binned in 5 μm increments relative to the glomerulus boundary. Dashed line shows the average distance between the central target location and the glomerulus edge. (**G**) The effect of laser power on onset PN spike rate (mean ± SEM), directly measured from PN recordings (n = 68; black) or inferred from detected EPSC times in KC recordings (n = 27 KCs, 108 synapses; red). Asterisks indicate significant differences between spike rates measured in PNs and KCs (p < 0.05, t-test with Holm-Bonferroni correction). **(H)** The relationship between onset spike rate and first spike latency (mean ± SEM) for control PN recordings (n = 68) and KC recordings (n = 27 KCs, 108 synapses). Asterisks indicate significant differences between PNs and KCs (p < 0.05, t-test with Holm-Bonferroni correction). Averaged across rates, KC latencies were delayed 10.3 ± 1.8 ms relative to PNs. **(I)** Rate-normalized ISIs were calculated for PNs (n = 68, SD = 0.12) and KCs (n = 108 PN-KC synapses, SD = 0.21) using the time-rescaling theorem, with all distributions having a mean of 1 by definition. A Poisson distribution with rates matched to PNs is shown for comparison (SD = 1.0). Plotted data include all ISIs across all trials.

To first identify which glomeruli were connected to a recorded KC, we mapped connectivity in current clamp by stimulating voxels tiling the antennal lobe (∼1,200 voxels per experiment; Figures 1C and S2). Glomeruli that evoked consistent KC depolarizations across trials were manually identified online, and the approximate center was selected for further investigation. However, temporal summation prevented reliable detection and measurement of excitatory postsynaptic potentials (EPSPs). For STP characterization, we therefore switched to voltage clamp where individual EPSCs could be resolved, and both spiking and feedback inhibition were suppressed (Figures 1D and S1).

Because our STP analyses required inferring PN spike times and measuring the associated EPSCs, we performed control recordings on PNs to validate our stimulation protocol and quantify the statistics of evoked spike trains (Figures 1E-1I). These recordings showed: (1) laser power reliably controlled PN spike rate, (2) inter-spike intervals (ISIs) were highly regular above ∼30 Hz, and (3) single-glomerulus spatial resolution was maintained for rates up to ∼60 Hz. However, sparse connectivity further improved the effective spatial resolution and enabled us to present stimuli up to ∼100 Hz, as our mapping empirically showed that pairs of connected glomeruli were rarely axially adjacent. Together, these measurements established a stimulation protocol that evoked sufficiently stereotyped PN responses to support PN spike time inference and quantification of the corresponding monosynaptic EPSC amplitudes.

For each identified synapse, we evoked PN spike trains using a range of laser powers and quantified the resulting EPSC amplitudes. Each trial consisted of two stimulation trains (500 and 250 ms) separated by a short delay (50-1,000 ms; Figures 1B and S2A), allowing us to characterize both synaptic modulation and the recovery rate toward baseline amplitudes. While response profiles were reproducible across trials for a given PN-KC synapse, substantial variation was observed across the population, ranging from strongly facilitating to strongly depressing (Figures 1D, S3, and S4). Thus, our experimental approach revealed diverse synaptic dynamics across a range of presynaptic firing rates for individual PN-KC connections.

### Tsodyks-Markram model provides a rate-independent description of heterogeneous synaptic dynamics

STP is inherently rate dependent such that the response at one spike rate provides only a snapshot of the underlying dynamics with limited predictive power for other presynaptic rates. Clustering trials by inferred PN spike rate highlighted this feature of STP, as EPSC amplitudes typically varied across input frequencies (Figures 2A, S3, and S4). This rate dependence can pose an analytical challenge for comparing synaptic dynamics across a population and determining its organization. More importantly, neural circuits respond to a wide range of spiking statistics, with naturalistic stimuli exhibiting substantial variability in spike rates and spike timing. Understanding how STP contributes to network-level computations therefore requires a more generalized framework for describing STP than single-rate measurements can offer.

**Figure 2.**
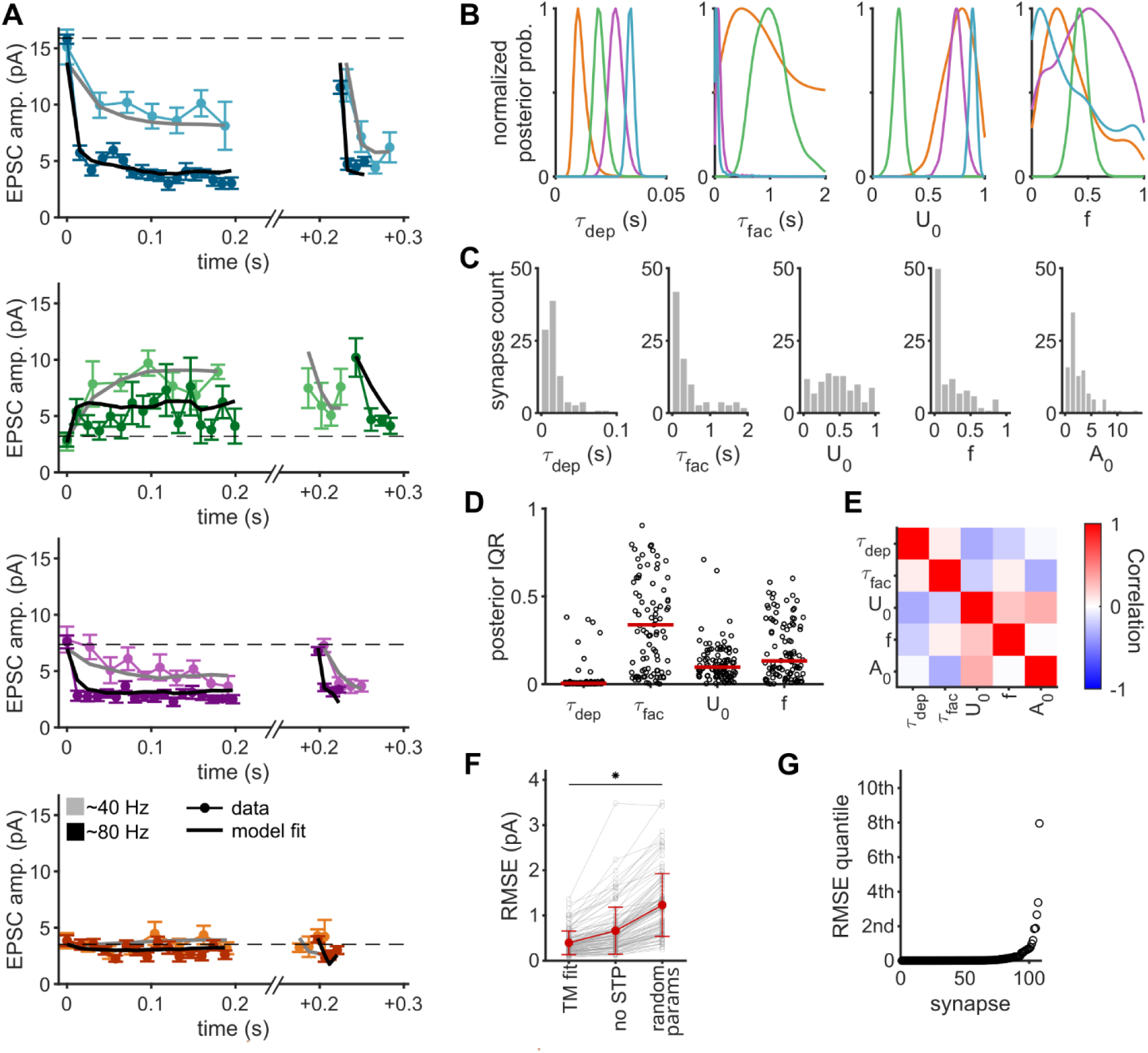
Tsodyks-Markram model provides a rate-independent quantification of heterogeneous STP dynamics. **(A)** EPSC amplitudes (mean ± SEM) and Tsodyks-Markram fits for the KC shown in Figure 1. Light and dark shades average across trials with inferred PN spike rates near 40 Hz and 80 Hz. Only the first 200 ms of the stimulus 1 response is shown for visual clarity. Stimulus 2 is plotted relative to the final EPSC of stimulus 1, with experimental limitations preventing precise control of the delays (STAR Methods). **(B)** Posterior probability distributions for Tsodyks-Markram parameters, fit using a published Bayesian algorithm^44^. Estimated parameter values were selected as the peak of the marginal posterior distributions. A_0_ was fit using least squares. **(C)** Distribution of estimated parameter values (n = 27 KCs, 108 synapses). **(D)** Widths of the marginal posterior distributions (25th-75th percentile), with red lines showing median values. **(E)** Correlation matrix of estimated parameter values. **(F)** RMSE between measured and predicted EPSC values. Fits were compared to a fixed-weight model (no STP, zero-slope line) and to randomly selected parameter combinations (averaged across 100,000 combinations). All groups were significantly different from each other (p < 0.001, t-tests with Holm-Bonferroni correction). **(G)** Quantile values for model fits compared to 100,000 randomly selected combinations of Tsodyks-Markram parameters.

To capture this rate-dependent behavior, we fit our measured EPSCs with the Tsodyks-Markram model of STP^40,41^ (Figures 2, S3, and S4; STAR Methods). This phenomenological model reproduces a wide range of STP phenotypes across different ISIs using five synapse-specific parameters that describe history-dependent changes in synaptic efficacy according to:

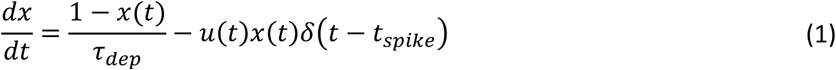

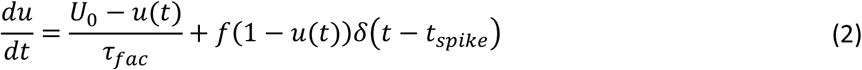

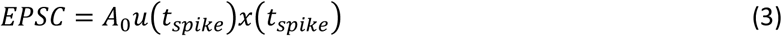

where x(t) is the proportion of synaptic vesicles available for release and u(t) is the release probability. The fixed parameters fit to each recorded synapse represent baseline EPSC amplitude (A_0_), timescale for recovery from depression (τ_dep_), timescale for decay of facilitation (τ_fac_), baseline release probability (U_0_), and facilitation strength (f).

Fitting the Tsodyks-Markram model using standard point-estimation methods can be highly sensitive to initial conditions and prone to local optima, so we instead used a previously published Bayesian algorithm that samples the joint posterior distribution using Markov chain Monte Carlo^44^. We selected the marginal posterior modes as parameter estimates for each synapse (Figure 2B). These marginal modes were less sensitive to small changes in the fitting procedure compared to the joint maximum *a posteriori* estimate while having a negligible effect on fit quality (ΔRMSE = 0.005 ± 0.009 pA; STAR Methods). All reported fits were robust to small changes in the fitting procedure (STAR Methods) and across independent fits using separate trials (Figure S5).

Marginal posteriors consistently showed clear unimodal peaks with concentrated probability densities (Figures 2D, S3E, and S4D). Estimated parameters accurately reproduced measured EPSC amplitudes (RMSE = 0.39 ± 0.26 pA), reducing error by 41% compared to fixed-weight control models (RMSE = 0.66 ± 0.52; p < 0.01, Figure 2F). Compared to 100,000 randomly generated parameter sets, selected estimates were the best-fitting parameter set for 26% of synapses and ranked within the top 0.5% for 90% (Figure 2G). Synapses with smaller relative improvements in RMSE typically had limited synaptic modulation (i.e. paired-pulse ratios near one). Average posterior density was moderately correlated with RMSE across randomly selected parameter sets (ρ = −0.55 ± 0.22, Spearman’s rank correlation), with lower posterior support associated with greater variability in RMSE (Figures S2–S4).

Parameter estimates across the population were heterogeneous and spanned a broad range of values (Figure 2C). Correlations between parameters were small, with the exception of moderate correlations between U_0_ and all other parameters (r ≈ ± 0.25, Figure 2E). Distributions for τ_dep_, f, and A_0_ were best fit by log-normal distributions, while τ_fac_ was best fit by a log-uniform distribution. U_0_ was instead best fit by a normal distribution, potentially reflecting additive contributions from the relative abundance of synaptic release proteins UNC13A and UNC13B^45,46^. These results therefore establish a common, rate-independent framework for describing heterogeneous synaptic dynamics, allowing us to examine how synaptic diversity shapes circuit function.

### A near-random organization of STP creates a heterogeneous set of temporal filters

To ask whether the observed STP heterogeneity had a structured organization across the circuit, we first performed hierarchical clustering on the estimated Tsodyks-Markram parameters (Figure 3A). Clustering identified three broad groups that roughly correspond to low-pass, high-pass, and band-pass temporal filters based on the predicted steady-state response to different presynaptic spike rates (Figure 3B). However, substantial and qualitatively continuous variation was observed within each cluster, suggesting each PN-KC connection expressed a unique signature of STP. Mapping cluster assignments onto the connectivity diagram of the recorded population revealed an intermingled pattern with no immediately evident organization based on KC or glomerulus identity (Figure 3C), suggesting a potentially independent assignment of STP phenotypes across the population.

**Figure 3.**
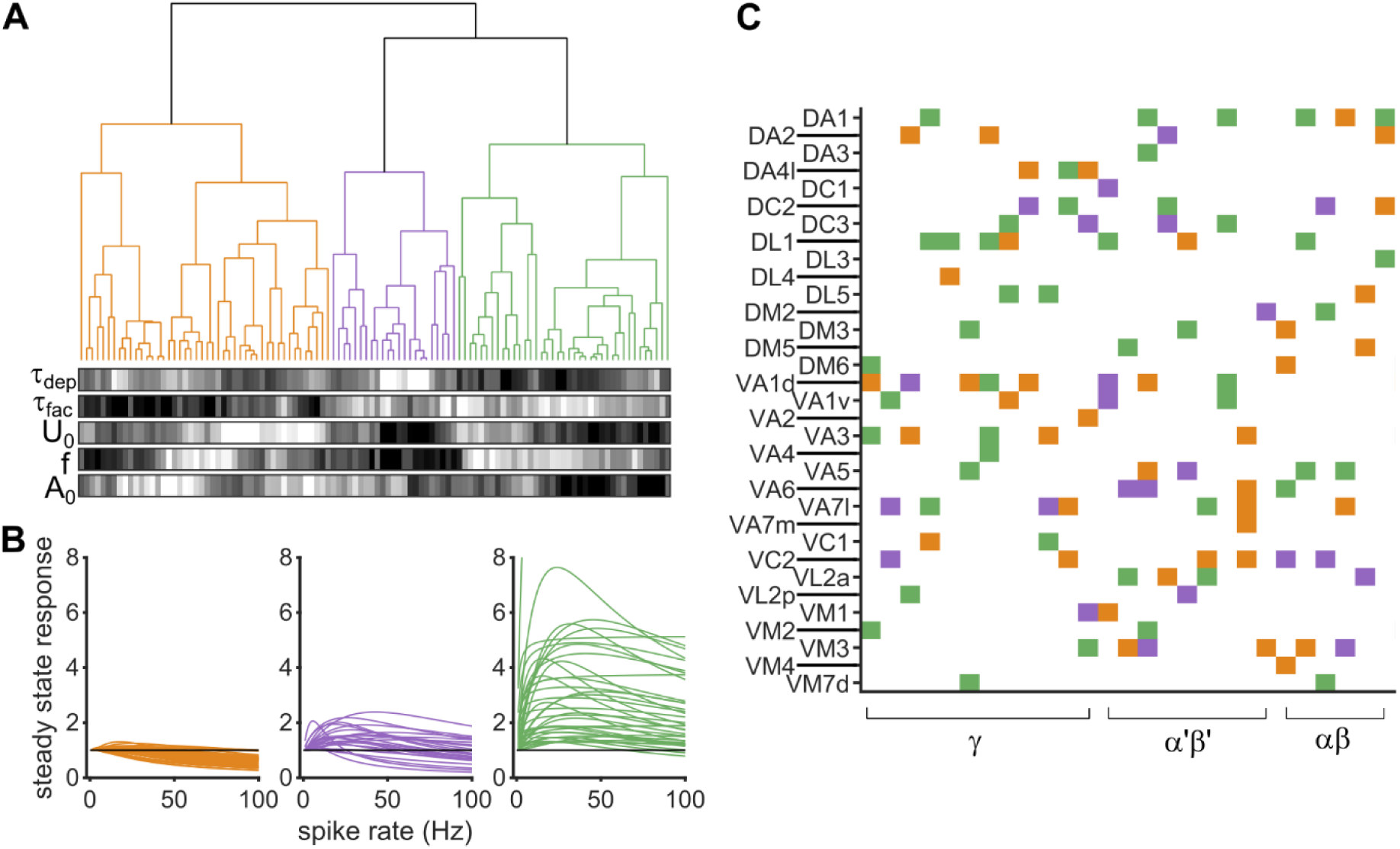
STP is heterogeneous across PN-KC synapses. **(A)** Hierarchical clustering of estimated Tsodyks-Markram parameters (n = 27 KCs, 108 synapses). Heat maps show the parameter estimates for each data point, scaled from minimum (black) to maximum (white). **(B)** Predicted steady-state responses for each synapse, simulated with spike trains from 1 to 100 Hz. Colors correspond to cluster identity in panel A. **(C)** Connectivity diagram of all recorded PN-KC pairs. Each row is a glomerulus and each column is a KC, sorted by subtype.

Because STP is predominantly a presynaptic phenomenon, we first evaluated whether phenotypes exhibited a structured organization based on glomerulus identity (Figure 4A). We computed all pairwise distances between the estimated Tsodyks-Markram parameters and segregated them into synapses from the same glomerulus and synapses from different glomeruli. Comparing these distributions showed no significant difference based on glomerulus identity (Figure 4B).

**Figure 4.**
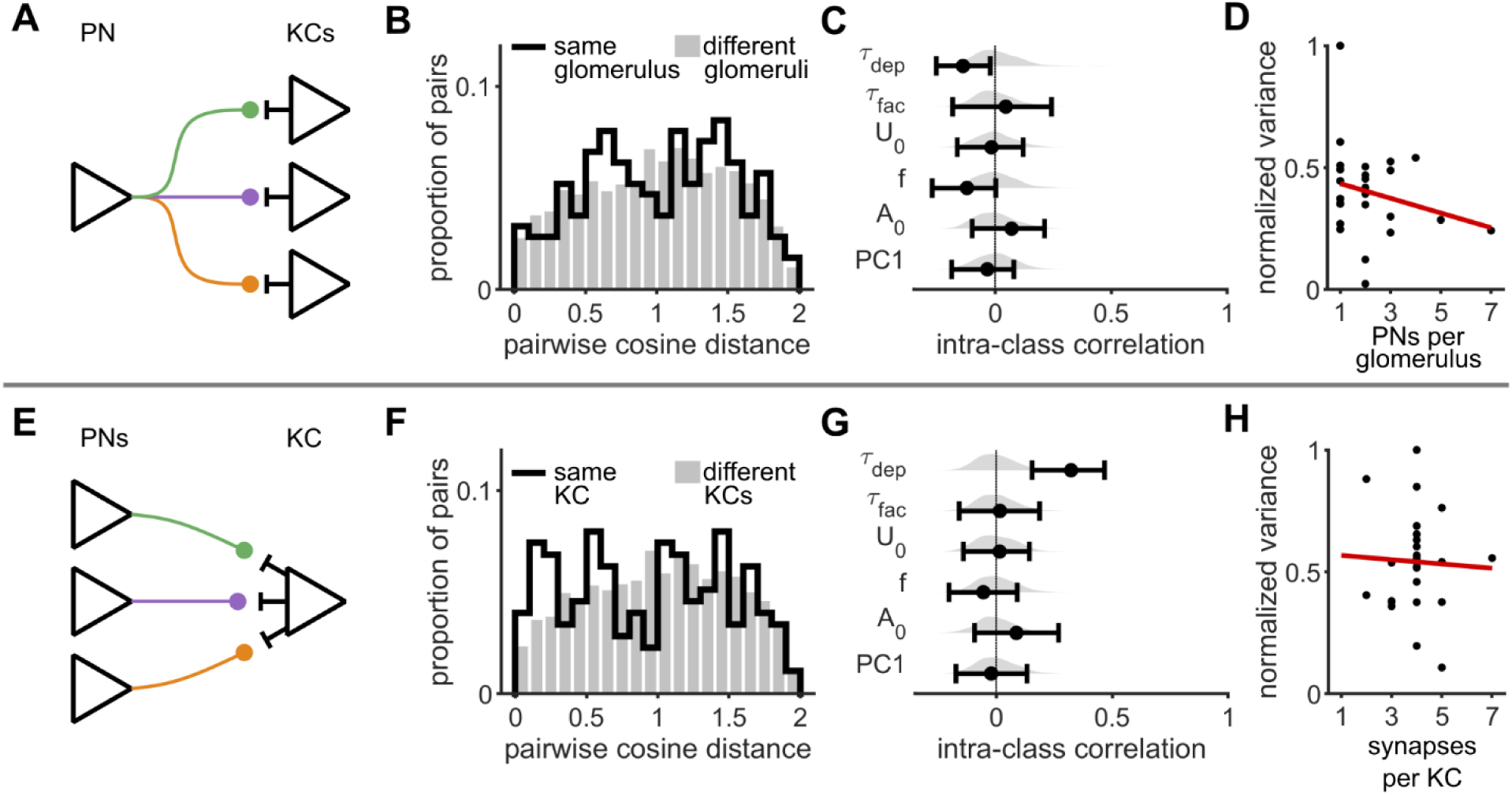
A near-random organization of STP across the circuit. **(A)** Schematic illustrating the potential organization of STP phenotypes around PN identity. **(B)** All pairwise distances between Tsodyks-Markram parameter estimates (n = 108), separated into pairs from the same glomerulus and pairs from different glomeruli (p = 0.55, permutation test with 10,000 shuffles). **(C)** Intraclass correlations measured for each fit parameter and the first principal component (36.8% explained). This metric measures how strongly parameters cluster based on glomerulus identity. Shaded histograms show the null distribution (10,000 shuffles), and error bars show the 95th percentile confidence interval (10,000 bootstraps). **(D)** Comparison of parameter variance relative to the number of PNs per glomerulus. Each point is a glomerulus, with the y-axis representing parameter variance (trace of covariance matrix) and the x-axis representing the number of PNs in the glomerulus (counted in hemibrain connectome). Red line shows the linear fit (R^2^ = 0.06, p = 0.25). **(E-H)** Same as A-D but analyzed based on KC identity. Distributions in panel F were significantly different (p = 0.04, Cohen’s d = 0.14, permutation test with 10,000 shuffles). Panel H plots the number of recorded synapses per KC (R^2^ = 0.01, p = 0.82).

We further tested for organization based on glomerulus identity by calculating intraclass correlations (ICCs), a metric designed to quantify group-level clustering in nested data (Figure 4C). While statistical tests of pairwise distances collapse the distributions into group means and often violate assumptions of independence, ICC instead uses the full spread of data to partition the variance and directly measures how much variance is attributable to glomerulus identity. ICCs for individual parameter estimates were often negative, indicative of greater within-glomerulus variability than expected by chance, but none were significantly different from 0, indicating that STP phenotypes did not cluster by glomerulus identity. Since ICC is necessarily a univariate measure, we performed principal component analysis (PCA) and calculated ICCs for each PC as a multivariate approximation, similarly finding no evidence for clustering based on glomerulus identity (Figures 4C and S6). Simulations matched to experimental sample sizes recovered defined ICCs with SD = 0.08, indicating that we can confidently reject the presence of moderate or strong clustering.

Our stimulation protocol provided single-glomerulus spatial resolution, but glomeruli have variable numbers of PNs. This could result in greater observed variability in glomeruli containing more PNs, potentially masking organization based on individual PN identity. We therefore quantified within-glomerulus variance and compared it to the number of PNs per glomerulus (Figure 4D). This comparison yielded a nonsignificant negative correlation, further supporting a conclusion that STP phenotypes were not organized around PN identity.

Postsynaptic cell identity can also influence STP phenotypes, so we next asked whether STP was organized by KC identity (Figure 4E). Comparing pairwise distances showed that PNs synapsing onto the same KC were more similar than expected by chance (Figure 4F), although the associated effect size was small (p = 0.04, Cohen’s d = 0.14). ICC analyses suggested τ_dep_ was primarily responsible for this small shift (ICC = 0.32), as all other parameters had ICCs indistinguishable from zero (Figures 4G and S6). Notably, these results represent an upper bound on KC-level clustering because PNs converging onto the same KC share potentially confounding sources of variability such as fly identity. The number of recorded PNs per KC did not influence these results (Figure 4H), and grouping synapses by KC subtype (γ, α’β’, αβ) similarly showed limited evidence for structured organization (Figure S6).

Together, these analyses revealed extensive heterogeneity in STP across PN–KC synapses with little evidence for a structured organization. STP instead appeared to be organized randomly and independently, resulting in an uncorrelated distribution of temporal filters across the circuit. Mirroring the pseudo-random anatomical connectivity important for pattern separation^33,34^, this unstructured organization raised the possibility that heterogeneous STP might similarly decorrelate temporal information into structured representations.

### Network models with heterogeneous STP reproduce KC population clock dynamics

We next asked how the observed heterogeneity in STP shapes emergent KC population dynamics using an experimentally constrained model of the mushroom body (STAR Methods and Table S1). KCs were modeled as 2,000 conductance-based, leaky integrate-and-fire neurons (LIF) that were randomly connected to 6 out of 50 PNs. PN odor response peristimulus time histograms (PSTHs) were parameterized to resemble realistic rate trajectories^39,47,48^, and synapses were randomly assigned Tsodyks-Markram parameters from a distribution matching the experimentally measured population. Network inhibition was tuned to match experimentally measured population sparseness^49–51^. Results were robust to a range of biophysical parameters and network architectures (Figure S7A).

We first compared the population dynamics of our model with previously published KC odor responses that consisted of 82 α‘β‘ KCs responding to a 10 s odor stimulus^37^. We simulated matching 10 s odor presentations and evaluated KC spike rates across repeated presentations of the same odor. Ordering KC PSTHs by peak latency revealed a complete but non-uniform tiling of stimulus time (Figure 5A). We quantified temporal coverage using the participation ratio (PR), where higher values indicate a more uniform distribution of peak latencies. Temporal coverage was higher for models (PR = 0.60 ± 0.02) compared to published data (PR = 0.45; Figure 5C), although this difference can largely be explained by a smaller number of KCs in the experimental dataset (sample size matched PR = 0.53 ± 0.03). We further asked whether individual KCs exhibited consistent peak latencies across different odors (STAR Methods). Only 1.9 ± 0.5% of KCs exhibited a significant latency preference across odors, suggesting that temporal structure emerges at the population level rather than through temporal specialization of individual KCs.

**Figure 5.**
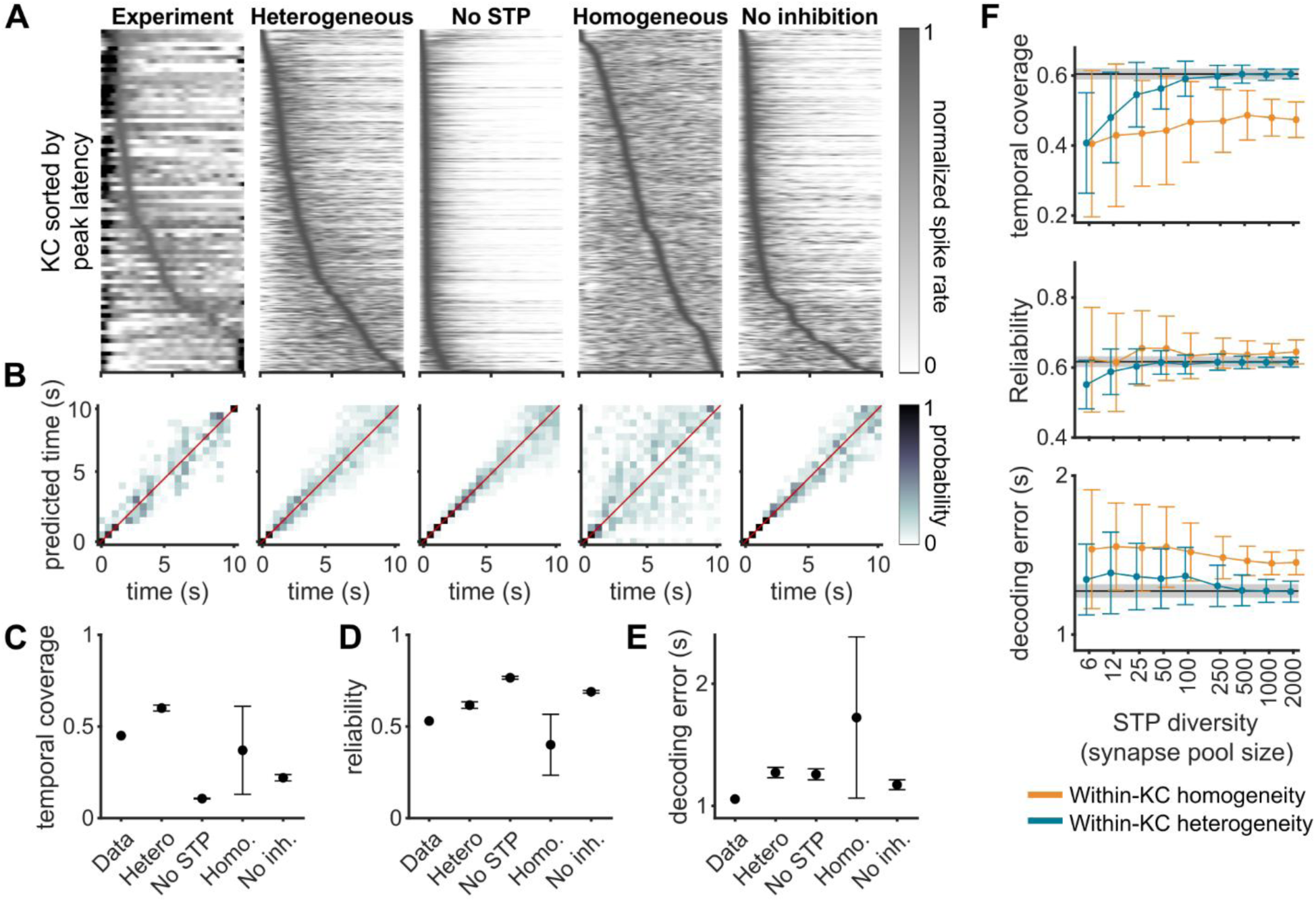
Network models require heterogeneous STP to reproduce *in vivo* population clock dynamics. **(A)** KC odor response PSTHs for previously published *in vivo* odor responses^37^ (left, n = 82) and 4 representative network simulations with different synaptic dynamics. Each row is a KC odor response normalized by its maximum spike rate and sorted by peak-response latency. Normalization and sorting of the experimental data excluded the strong onset response (0-1 s, saturated black), following the original publication. For heterogeneous models, each synapse was randomly assigned experimentally constrained Tsodyks-Markram parameters. For no STP models, all synapses were assigned a fixed weight. For homogeneous models, all synapses were assigned identical parameters. No inhibition models have heterogeneous STP and reduced excitatory gain. Each example network has between 230 and 280 KCs responding to the same simulated odor. **(B)** Confusion matrices show Bayesian decoding of stimulus time for the example networks in panel A. Color indicates the posterior probability assigned to each time bin (500 ms width). **(C-E)** Quantification of temporal structure across network replicates (mean ± SD, n = 1, 20, 20, 200, and 20, with each network averaged across 50 unique odors). **(C)** Temporal coverage measured using the participation ratio, which indicates how uniformly peak latency estimates tile temporal space. **(D)** Reliability measured as Spearman’s rank correlation between peak latency estimates obtained from independently simulated trials. **(E)** Decoding error was measured as the RMSE between stimulus time and predicted time, averaged across the full stimulus duration. **(F)** STP heterogeneity was systematically varied by generating pools of potential STP phenotypes of various sizes (x-axis). For within-KC heterogeneity models, all synapses were independently assigned a phenotype from the available pool. For within-KC homogeneity models, each KC sampled a phenotype that was assigned to all 6 of its PN inputs. Each data point represents 50 different networks (mean ± SD), each generated with a unique pool. Black lines show results for the heterogeneous models in panels C-E. **(A-E)** Experimental data for *in vivo* KC odor responses were reproduced from Kropf et al. (2026)^37^, used under a CC BY 4.0 license.

Because KCs spiked throughout the stimulus, we next evaluated response sparseness and reliability. Sparseness had similar means but a narrower distribution for models compared to experimental data (Figure S7B), potentially reflecting intrinsic heterogeneity not accounted for in models. To evaluate reliability, we compared peak latency measurements across independent trial subsets for each network-odor pair (Figure 5D). Response reliability was moderate for both data (ρ = 0.53, Spearman’s rank correlation) and models (ρ = 0.62 ± 0.02), suggesting a coarse rather than precise representation of time.

Temporal coverage and reliability describe the population dynamics but do not directly quantify the information they carry about stimulus time. We therefore implemented a Bayesian time decoder that infers stimulus time based on the full population response binned into 500 ms intervals across the 10 s odor stimulus^37^ (Figure 5B). Decoding accuracy was similar between published data (RMSE = 1.06 s) and models (RMSE = 1.27 ± 0.04 s; Figure 5E), corresponding to 2-3 time bins of temporal uncertainty. We used average RMSE as a summary metric but note that error magnitudes were not uniform across time, increasing linearly at similar rates for data and models (Figures 5B and S7C). Together, these three analyses showed that our experimentally constrained model reproduces the major features of KC odor response dynamics, providing a framework for exploring the circuit mechanisms underlying this temporal representation.

### Population clock dynamics require heterogeneous STP

While our full model accurately reproduced experimental observations, it remained unclear whether heterogeneous STP was necessary and sufficient for the emergence of a population clock. To address this question, we altered STP organization and evaluated changes in KC responses. We began by fully removing STP from the model and assigning a fixed weight to each synapse (STAR Methods). These networks exhibited strong onset transients and limited sustained activity, resulting in low temporal coverage (PR = 0.11 ± 0.002; Figures 5A and 5C). In contrast, temporal decoding remained comparable to the models with STP (RMSE = 1.26 ± 0.05 s; Figures 5B and 5E). This result likely reflects a population-level rate code where the population PSTH provides sufficient information for the Bayesian algorithm to infer time even in the absence of an explicit population clock.

Fixed-weight models demonstrated that STP is required for population clock dynamics but did not establish whether a heterogeneous organization is needed. We therefore generated homogeneous networks where all synapses were assigned identical STP properties. Across homogeneous networks we observed an inverse correlation between temporal coverage and decoding accuracy (r = −0.78, Figure S7F), which furthermore corresponded to the paired-pulse ratio of the network’s identical synapses. Networks with a paired-pulse ratio near one behaved like a fixed-weight network with low temporal coverage and good decoding. In contrast, both facilitating and depressing networks with greater synaptic modulation typically had extended temporal coverage but poor decoding that often approached chance levels (Figure S7). These results suggest STP can generate temporal structure, but without heterogeneity this structure fails to produce the combination of temporal coverage and accurate decoding inherent to a population clock.

The apparent requirement for STP heterogeneity next raised the question of how much diversity is needed to shape KC dynamics into a robust population clock (Figure 5F). The full model independently assigned a unique phenotype to each synapse. To systematically vary the degree of heterogeneity, we generated a limited pool of available phenotypes (range 6 to 2,000) that each synapse sampled from. We tested two different sampling rules. In the first, each synapse randomly selected an STP phenotype (within-KC heterogeneity), matching the selection process of the full model. In the second, each KC sampled an STP phenotype that was shared across all of its convergent inputs (within-KC homogeneity). For within-KC heterogeneous networks, performance metrics converged to the mean dynamics of the full model with pool sizes between 250 and 500, potentially reflecting the small number of active KCs per odor (Figure 5F). Variance converged more slowly, requiring a pool size near 1,000. Within-KC homogeneous networks failed to reproduce these dynamics even when every KC was assigned a unique STP phenotype (i.e. pool size = 2,000; Figure 5F). These results indicate that heterogeneity among convergent synaptic inputs rather than population-level diversity alone is required for population clock dynamics.

Finally, we asked whether heterogeneous STP alone could reproduce the observed population dynamics or whether additional circuit mechanisms were required. Removing inhibition from our models doubled the number of active KCs per odor and produced strong onset transients resembling fixed-weight networks (data not shown). Reducing excitatory gain to restore sparse activity partially recovered population clock dynamics, but peak latencies remained biased towards stimulus onset (PR = 0.22 ± 0.02; Figures 5A and 5C). In contrast, reliability (ρ = 0.69 ± 0.01; Figure 5D) and decoding accuracy (RMSE = 1.17 ± 0.04 s; Figure 5E) remained comparable to models with inhibition. Thus, inhibition appears to play an important but secondary role by regulating excitatory drive and distributing peak latencies more evenly across stimulus time, consistent with recent work in the cerebellum^27^.

Together, these results establish STP heterogeneity as the primary mechanistic source of KC population clock dynamics, with inhibition refining and extending the distribution of the peak latencies. The resulting representations depend not only on a heterogeneous organization, but also on its distribution among synapses converging onto the same cell.

### Heterogeneous STP reformats latent temporal information into an MBON-decodable population clock

Across models, temporal decoding was remarkably similar despite large differences in temporal coverage. However, only the full model incorporating heterogeneous STP and inhibition reproduced the combination of both broad temporal coverage and accurate decoding. The relative invariance in decoding accuracy suggested that temporal information may originate upstream of the KC population. Consistent with this idea, temporal decoding from PN spike trains was more accurate than the KC population (RMSE = 0.75 ± 0.17 s). Fisher information analyses provided an alternative estimate of temporal uncertainty, showing that PN PSTHs can theoretically resolve time with a precision greater than the temporal resolution of the Bayesian decoder (Figure 6A).

**Figure 6.**
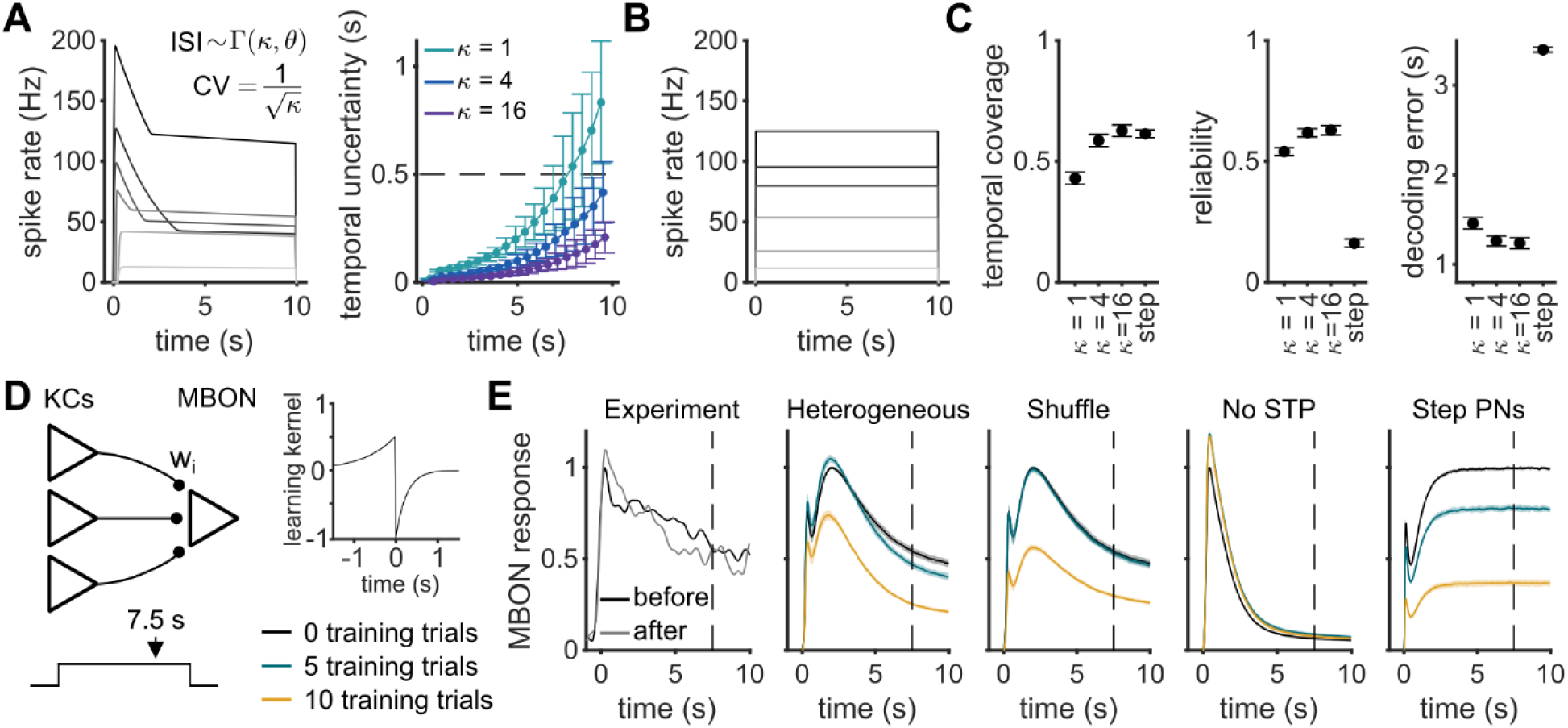
Population clock dynamics provide an MBON-decodable representation of latent PN temporal information. **(A)** Each line in the left panel is an example PSTH for a simulated PN odor response. The right shows temporal information in PN rate trajectories, quantified using the Cramér–Rao bound derived from population Fisher information (mean ± SD, n = 50 simulated odors). Dashed line indicates the temporal resolution for all PN and KC analyses. Equation shows how spike times were sampled for each simulated trial, with θ representing spike rate and ISI variability controlled by the shape parameter κ. **(B)** Example PN PSTHs designed as step functions to remove any information about stimulus time. **(C)** Quantification of temporal structure for KC populations simulated with PN inputs containing different amounts of temporal information (mean ± SD, n = 20). **(D)** MBONs were modeled as rate-based leaky integrators receiving input from all KCs, with an unconditioned stimulus occurring 7.5 s after odor onset (left). Synaptic weights were updated according to an STDP learning rule (right). **(E)** MBON responses before and after learning for previously published data (left) and different network manipulations (mean ± SD, n = 20). Shuffle denotes heterogeneous models where KC identity was randomly permuted between training and test trials. Experimental MBON recordings were reproduced from Kropf et al. (2026)^37^, used under a CC BY 4.0 license.

To test whether KCs inherit temporal information from PNs, we manipulated the temporal structure of PN odor responses (Figures 6A and 6B). PN spike trains were drawn from a gamma distribution, where increasing the shape parameter κ decreases both ISI variability and temporal uncertainty (Figure 6A). Increasing κ from 1 (Poisson) to 4 significantly increased KC temporal coverage, reliability, and decoding accuracy (Figure 6C). A further increase to κ = 16 produced only negligible improvements in these metrics, consistent with temporal uncertainty in both conditions falling below the temporal resolution of decoding analyses.

For a larger perturbation to PN temporal information, we generated new PN odor responses that had step-function PSTHs intentionally designed to remove all information about stimulus time (Figure 6B). Networks simulated with these odors produced comparable temporal coverage to standard PN models (PR = 0.61 ± 0.02) but had significantly lower reliability (ρ = 0.16 ± 0.02) and decoding accuracies approaching chance levels (RMSE = 3.40 ± 0.03 s; Figure 6C). Thus, KCs inherit temporal information embedded in PN rate trajectories, with heterogeneous STP functioning to organize this information into a structured, population-level representation of time.

Because most network types could accurately decode inherited temporal information, it remained unclear whether the STP-mediated transformation into a population clock served a functional purpose. Accurate decoding by an ideal observer does not necessarily imply that information is accessible to a biologically constrained decoder, so we next simulated an MBON as a rate-based LIF neuron (Figure 6). MBONs receive convergent input from all KCs of a given subtype, with dopamine-mediated LTD at the KC-MBON synapse playing a key role in olfactory associative learning. MBON spike rates then provide an accurate readout of learned odor valence^35,52^. Previous work showed that repeated reinforcement 7.5 s after odor onset led to behavior and MBON plasticity that were both temporally specific, akin to classical eyeblink conditioning^37^. Our MBON simulation reproduced this paradigm, using an experimentally motivated spike-timing-dependent plasticity (STDP) rule to update KC-MBON weights following an unconditioned stimulus^36^.

Only networks with heterogeneous STP could reproduce temporally specific MBON plasticity. This plasticity included LTD that was coarsely aligned with the reinforcement signal along with a slight potentiation of the onset response (Figure 6E). Stronger reinforcement (i.e. more training trials or higher learning rate) resulted in a more global depression resembling the LTD more typically reported in MBONs^35^ (Figures 6E and S8). Shuffling KC identity between training and test trials abolished temporally specific learning, but global learning was retained with stronger reinforcement (Figure 6E). Networks with no STP could not learn, while step-function PNs resulted in only global MBON depression. Earlier or distributed reinforcement times similarly produced a more global depression, consistent with previous reports (Figure S8).

Overall, these results clarify the computational function of heterogeneous STP as a distributed set of temporal filters that transforms temporal information already present in PN activity into a structured representation of time. This reformatting of temporal information plays a central role in supporting MBON plasticity that directly drives learned behaviors^29,37^. Interestingly, while the KC population clock was necessary for downstream plasticity, this temporally structured representation was ephemeral, converting back into a rate code in MBONs.

### STP enhances pattern separation for similar odors

The mushroom body primarily functions as a pattern separation circuit where odor identity is encoded by the sparse set of odor-responsive KCs^33,34^. We asked whether heterogeneous STP additionally contributes to this computation by generating odors with defined similarities and comparing overlap in the set of active KCs (Figure 7A). At low odor correlations, networks with and without STP both effectively separated odor identity, although fixed-weight networks did perform better. Interestingly, this relationship reversed at high correlations (Figure 7B), with STP networks producing greater pattern separation than fixed-weight models. Networks with STP had a narrower range in the number of active KCs per odor that may contribute to this improved separation (Figure 7C), consistent with a role for STP in gain control^53^. However, the similarity-dependent effects suggest that gain control alone cannot fully explain these results. Instead, heterogeneous STP appears to enhance odor identity coding specifically when odors are similar and discrimination is difficult.

**Figure 7.**
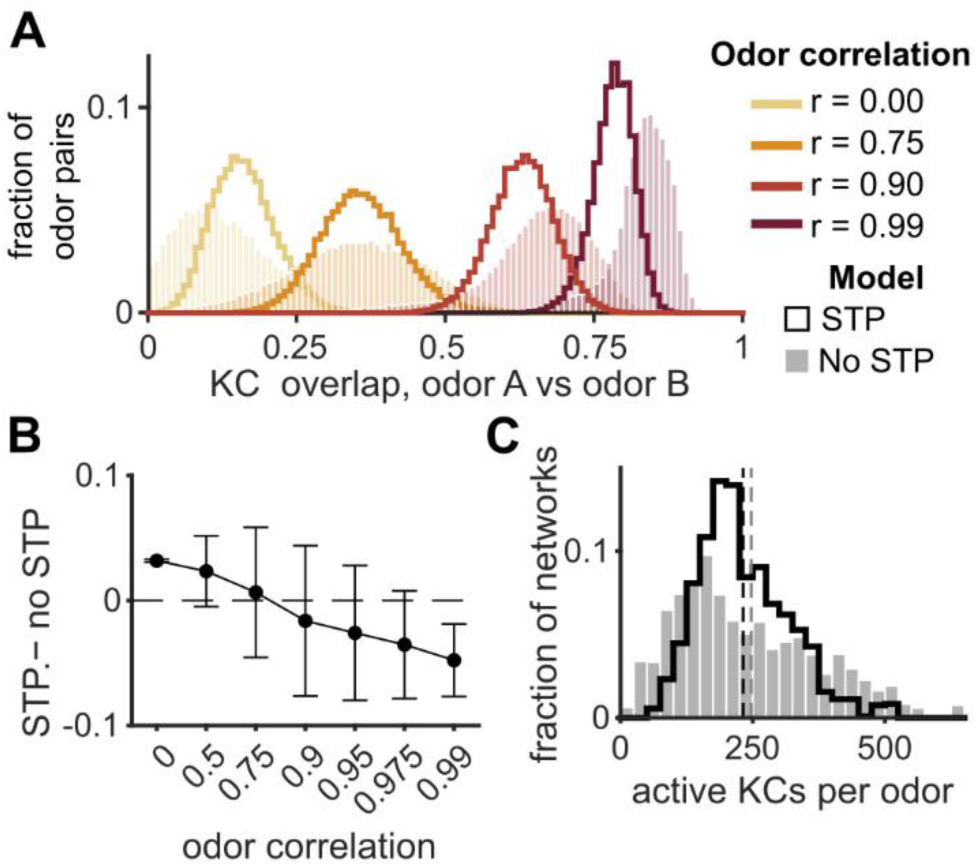
Heterogeneous STP enhances pattern separation for similar odors. **(A)** Pattern separation of odor identity was quantified as the overlap in active KCs (Dice–Sørensen coefficient) for a network (n = 5) responding to a pair of odors with a fixed PSTH correlation (10 odor families per correlation, 50 odors per family). Lines show models with heterogeneous STP and shading shows models with fixed-weight synapses. **(B)** Difference in average overlap between networks with and without STP (mean ± SD, n = 10 odor families). Note that the x-axis is scaled categorically for visual clarity. **(C)** Histogram shows the number of active KCs per odor. Plotted data include 20 networks responding to 50 uncorrelated odors.

## DISCUSSION

Transforming afferent information into formats accessible to downstream circuits is a key function of neural circuits. While ideal observers can be designed using unrestricted operations to optimally extract information, biophysical constraints limit the computations available to a biological decoder. Here, we showed that a heterogeneous organization of STP bridges this gap by providing a distributed circuit mechanism that reformats latent, rate-based temporal information into an explicit, MBON-interpretable population clock. This functional heterogeneity creates a temporal code largely orthogonal to anatomy-based pattern separation, though modest improvements to odor identity coding suggest some computational overlap. Together, these results demonstrate how biophysical heterogeneity can generate computational complexity without requiring a correspondingly complex circuit architecture.

Our results first identify heterogeneity as a distributed computational resource, linking computational building blocks like STP to circuit-level function. The experimentally observed heterogeneous organization comprised a qualitatively continuous distribution of temporal filters, potentially serving as a temporal basis set capable of decorrelating temporally complex PN inputs^23,27^. In contrast, homogeneous networks often failed to preserve existing temporal information, suggesting a common filter provides only limited temporal bandwidth. This differs from cortical models, where incorporating homogeneous STP can improve temporal coding^54–56^. The recurrent architecture of these models may provide an anatomical basis for temporal coding that any distribution of STP can enhance. In comparison, the predominantly feedforward anatomy of the mushroom body provided a tractable system for isolating and examining how STP and its organization help shape network dynamics.

The population clock generated by heterogeneous STP was an emergent phenomenon rather than a property of KCs individually tuned to different stimulus times. Neurons across sensory and motor systems can be selectively tuned to various temporal features^57^, and tuned STP offers one underlying mechanism^58–61^. However, model KCs did not have consistent peak latencies across different odors, in agreement with *in vivo* odor responses^37^, and suggesting a complex interaction between STP, inhibition, and PN spiking statistics. Network performance required heterogeneity among synapses onto the same KC, indicating that this temporal representation likely emerges from KCs integrating filtered PN input across local, independently functioning synaptic computations. The substantial variation in PN spike rates across odors could make this diversity of synaptic filters useful by allowing KCs to extract temporal structure that differs across stimuli^13^.

We observed a near-random organization of STP, but models reached asymptotic dynamics with substantially less heterogeneity. One explanation for this discrepancy could be that random assignment represents a low-cost developmental strategy. Full convergence of both means and variances required at least 1,000 unique phenotypes, far surpassing the population size of PNs (∼150 across 51 glomeruli). Second, additional heterogeneity could provide a buffer for trial-to-trial variability in PN spiking or contribute to different computations such as dynamically tracking an odor plume. Third, model fits may overestimate the true functional dimensionality, with fine-scale differences between synapses reflecting EPSC noise or uncertainty in parameter estimation. However, the persistence of substantial variability at intermediate levels of the dendrogram suggests the broader organization of STP diversity reflects biologically relevant heterogeneity.

Time is an important dimension for most behaviors, with population clocks providing one form of temporal representation^62^. Circuits can either passively inherit or actively generate temporal information, and the boundary often corresponds to sensory and motor systems, respectively^57^. Consistent with STP functioning as a passive temporal filter^23,63^, KCs could not create time *de novo* but instead reformatted information embedded in PN rate trajectories. KCs retained most of the available information even though the resulting population clock provided only a coarse tiling of temporal space. This level of temporal structure may flexibly balance precision and generalization^64–66^, enabling temporally aligned learning while allowing variable delays or stronger reinforcement to converge towards a global learning signal.

Population clocks were originally proposed to support spatiotemporal processing in the cerebellum^67^, providing an early framework for the temporal learning studied here in the anatomically similar mushroom body. In the vestibulocerebellum, three distinct populations of mossy fibers converge onto the same granule cells to support computations related to self-motion^68^. Recent work has demonstrated that STP biases matched to the statistics of these populations can support a short-timescale population clock^9,27^. With PNs typically considered as a single homogeneous class, our work builds on these results to show that incorporating fine-grained, within-class variability can further expand the computational repertoire of a circuit. This additional source of heterogeneity enabled the KC population to encode a consistent representation of time in response to diverse odors with variable temporal statistics. The KC clock also operates on a substantially longer timescale, with our results pointing towards inhibition as a key factor underlying this temporal extension.

Heterogeneity is an inherent feature of many complex systems, both within and outside biology, that can make fundamental contributions to system stability, adaptability, and overall performance^69–72^. In neural circuits, these contributions introduce an additional dimension to network computation, with recent work showing how variability in static biophysical properties such as spike thresholds and membrane time constants can diversify circuit responses^14–18^. Our work extends this framework to synaptic dynamics, showing circuit-level effects of heterogeneous STP that are consistent with broader principles of complex system function^69^. Overall, our results show how network-level dynamics can be built by and emerge from computational building blocks distributed across a neural circuit.

## Limitations

The Tsodyks-Markram model provided a tractable description of STP dynamics but abstracts much of the underlying physiology into a small set of phenomenological parameters. The model assumes a linear relationship between synaptic efficacy and its underlying state variables, while we did observe qualitatively non-linear dynamics that the model cannot describe. Given noisy data and finite sample sizes, fitting this model to data can also produce correlated and compensatory parameter estimates that could limit the resolution of comparisons between closely related dynamics, although our Bayesian approach reduced this risk. Finally, model parameters cannot be directly mapped to the underlying molecular machinery. Nevertheless, each parameter has a biological interpretation, and observing the highest variability in U_0_ and τ_fac_ suggests that synaptic calcium dynamics may be an important target for future work.

The mushroom body is a predominantly feedforward circuit with PNs comprising a single class of biophysically homogeneous neurons. This relative simplicity allowed us to isolate within-class STP heterogeneity and its contribution to circuit computations. However, cortical circuits, as well as other pattern separation circuits like the cerebellum and hippocampus^34^, introduce substantially more diversity in cell types, connectivity, and computations. Because the specific effects of heterogeneity can vary with cell type and circuit^14^, it will be important to determine how the STP diversity observed here scales and integrates into these increasingly complex systems.

## Acknowledgements

We would like to thank Isaiah Asbed and Jeff Johnston for comments on an earlier version of this manuscript. We would also like to thank Joel Greenwood and Omar Mano from the Yale Neurotechnology Core. This work was supported by NIH grant R01 NS116584, the Richard and Susan Smith Family Award for Excellence in Biomedical Research, the Klingenstein-Simons Fellowship Award in Neuroscience, and an innovative research award from the Kavli Institute for Neuroscience at Yale University to J.M.J. N.P.B. was supported by a fellowship from the Swartz Center for Theoretical Neuroscience.

## Author contributions

N.P.B. and J.M.J. designed the study. N.P.B. performed electrophysiology experiments, generated the model, analyzed the data and wrote the original manuscript. N.P.B. and J.M.J. revised the final manuscript. J.M.J. supervised the study.

## Declaration of generative AI and AI-assisted technologies in the writing process

The authors used ChatGPT 5.6 (OpenAI) to improve the manuscript’s readability and grammar. The authors reviewed and edited all output to ensure accuracy and take full responsibility for the content.

## RESOURCE AVAILABILITY

### Lead contact

Further information and requests for resources and reagents should be directed to and will be fulfilled by the lead contact James M. Jeanne.

### Materials availability

This study did not generate unique reagents.

### Data and code availability

- All reported data have been deposited at Zenodo and will be publicly available at the date of publication.
- All original code needed to reproduce reported analyses and network simulations have been deposited at Zenodo and will be publicly available at the date of publication.
- Any additional information required to reanalyze the data reported in this study is available from the lead contact upon request.

## STAR METHODS

### KEY RESOURCES TABLE

| REAGENT or RESOURCE | SOURCE | IDENTIFIER |
|---|---|---|
| Deposited data |  |  |
| Electrophysiology data | This study | 10.5281/zenodo.21285353 |
| Experimental models: Organisms/strains |  |  |
| *D. melanogaster*: GH146-Gal4 | BDSC | 30026 |
| *D. melanogaster*: UAS-CsChrimson::mVenus | BDSC | 55136 |
| Software and algorithms |  |  |
| MATLAB | MathWorks | mathworks.com |
| Python | Python Software Foundation | python.org |
| ScanImage v.2022.0 | Vidrio Technologies | Pologruto et al.<sup>73</sup> |
| Tsodyks-Markram model fitting algorithm | Costa et al.<sup>44</sup> | 10.3389/fncom.2013.00075 |
| LIF network model | This study | 10.5281/zenodo.21285353 |
| Custom MATLAB scripts | This study | 10.5281/zenodo.21285353 |

## EXPERIMENTAL MODEL AND STUDY PARTICIPANT DETAILS

Flies were raised on a standard cornmeal agar medium with rehydrated potato flakes (Carolina Biological Supply) and yeast on a 12 hour light, 12 hour dark light cycle at 25°C. All experiments were performed on 4-5 day old female flies with the genotype GH146-Gal4/+;UAS-CsChrimson::mVenus. Original stocks were obtained from the Bloomington Drosophila Stock Center.

## METHOD DETAILS

### Electrophysiology

Electrophysiology experiments were performed in an *ex vivo* preparation. The brain was dissected from the head capsule of cold-anesthetized flies and placed in a Sylgard-coated dish. The brain was positioned with the dorsal-ventral axis of the brain approximately vertical to allow both electrode access to posterior KCs and optical access to the anterior antennal lobe. This orientation was supported by a small Sylgard wedge and secured with fine tungsten wire. The dish was continuously superfused with fresh external saline at room temperature (20-22°C). External saline contained (in mM): 103 NaCl, 3 KCl, 5 TES, 8 trehalose dihydrate, 10 glucose, 26 NaHCO_3_, 1 NaH_2_PO_4_, 4 MgCl_2_, and 1.5 CaCl_2_ with osmolarity adjusted to 274-276 mOsm. The saline was bubbled with 95% O_2_ and 5% CO_2_ to reach an equilibrium pH of 7.3.

Whole-cell patch-clamp recordings were performed on both PNs and KCs. PNs with mVenus expression were randomly targeted for recording and KCs were randomly targeted with cell type determined *post hoc*. The internal pipette solution contained (in mM): 140 potassium aspartate, 10 HEPES, 1 EGTA, 1 KCl, 4 MgATP, and 0.5 Na3GTP, with osmolarity adjusted to 263 mOsm and pH 7.2. For KC recordings, 13 mM biocytin hydrazide was added for *post hoc* visualization on a confocal microscope, and PN recordings instead included 50 mM Alexa Fluor 568 for visualization of glomerulus boundaries under 2-photon illumination at 775 nm. Patch pipettes were pulled from borosilicate glass (1.5 mm outer diameter, 0.86 mm inner diameter) and fire-polished to a resistance of 6-8 MΩ for PNs and 14-17 MΩ for KCs. Recordings were collected using a Multiclamp 700b amplifier (Molecular Devices) with a CV-7B headstage (5 GΩ feedback resistor).

Current-clamp recordings were performed on both PNs and KCs. Recorded voltages were filtered online at 4 kHz and digitized at 10 kHz. Cells requiring somatic current injections greater than 40 pA (PNs) and 10 pA (KCs) to evoke spikes were discarded. Membrane potential was held near −55 mV, and cells requiring less than −15 pA (PNs) and −3 pA (KCs) to reach this potential were discarded. Input resistance was measured using 1 s current injections of −10 pA (PNs) and −2.5 pA (KCs), and time constants were estimated by fitting a single exponential to these current steps.

Voltage-clamp recordings were performed on KCs held at −65 mV. Series resistance was monitored but not compensated using −10 mV voltage steps and was typically on the scale of 2-5% of KC input resistance. Experiments were terminated if series resistance drifted by more than 20% of the original measurement. Recorded currents were filtered online at 1 kHz and digitized at 10 kHz. For analysis of EPSCs, traces were further filtered at 250 Hz using a two-pole Butterworth filter. Because EPSCs were well fit by a synaptic alpha function where τ = 1.8 ± 0.1 ms, this filtering is expected to decrease measured EPSC amplitudes by only ∼2.5% while significantly improving EPSC resolution.

Pharmacological blockade of inhibition was not included in our *ex vivo* preparation. Initial efforts to block inhibition with low concentrations of picrotoxin (2.5 μM) resulted in synchronous oscillations (1-5 Hz) of PN spiking (>100 Hz) that precluded STP analysis (Figure S1A). Intersectional genetic approaches to block inhibition were similarly precluded by technical limitations related to expression instability and driver silencing inherent to multi-transgene stocks that include GH146-Gal4. We therefore characterized inhibitory currents to evaluate the potential contribution of inhibition, obtaining results consistent with previous studies^50,74^ (Figure S1). Voltage clamp suppressed both spiking and inhibition except for some α’β’ cells. Trials with unclamped spikes or prominent inhibition were excluded from analysis. While GABAergic contamination below detection limits cannot be fully excluded, this characterization indicated that reported results predominantly reflect monosynaptic excitation with limited influence from unblocked inhibition.

### 2-photon optogenetic stimulation

Experiments were performed on a Scientifica Hyperscope with a Chameleon Ultra II Ti:Sapphire laser (Coherent), a 20x water immersion objective (XLUMPlanFL N, NA 1.0, Olympus), and a piezoelectric objective positioner (Physik Instrumente). The microscope was controlled using ScanImage version 2022.0 (Vidrio Technologies). The laser was tuned to 920 nm for antennal lobe imaging of mVenus expression, 1,040 nm for multiphoton optogenetic stimulation, and 775 nm for imaging Alexa Fluor 568 fills after PN recordings.

After establishing a whole-cell recording, we first acquired a reference image of the antennal lobe. This image consisted of 80 optical slices in 1 μm increments with an x-y resolution of 5.4 pixels/μm. The brain position exhibited slow drift during experiments, typically on the scale of 10-25 μm/hour. To compensate for this movement, we paused the experiment every 5-8 minutes and acquired a test image at a depth of 25 μm. We performed a 2D cross-correlation between this test image and each optical slice from the original reference image. We then identified the position that maximized the correlation and used this estimate to adjust stimulation coordinates and maintain spatial alignment. Experiments were terminated if larger-scale movement or nonlinear rotations of the brain prevented reliable image registration and accurate tracking of stimulation locations, all of which were typically associated with a degradation of recording quality.

We optogenetically evoked PN spike trains by stimulating spatially restricted voxels using the ScanImage Photostimulation Module^73^. Stimulation targeted 4 μm × 4 μm regions and was delivered using the built-in sinesquare function, which traversed the voxel every 10 ms. We used a pulsed stimulation protocol with 50 ms periods and a 40% duty cycle, except for the first period of each stimulus which had a 100% duty cycle. Stimuli also included spatial jitter such that each period shifted the center of the pattern to a location 2 μm from a central focus point. This parameterization of our 2-photon stimulation protocol was empirically optimized to maximize response reliability and minimize depolarization block using preliminary recordings from PNs.

To map PN-KC connectivity, we stimulated voxels tiling the antennal lobe while recording from a KC in current clamp. Voxels were spaced in a 6 μm grid and presented in a random order, typically resulting in 1,100-1,300 voxels per experiment. Stimulations were 250 ms in duration with a 250 ms inter-stimulus interval and presented at a power of 1.8 mW. These data were used strictly for connectivity mapping rather than STP quantification, so we prioritized presenting rapid, strong stimuli to facilitate connection identification. We manually identified connected PNs online by visually assessing which glomeruli evoked reliable KC depolarizations. We selected the approximate center of each connected glomerulus for additional stimulation and presented test stimuli to verify connectivity and determine the minimum laser power needed to evoke a detectable response, which was typically in the range of 0.2-0.5 mW. We discarded KCs in rare cases where two connected glomeruli were closely aligned along the axial dimension and single-synapse spatial resolution could not be ensured.

After identifying connected glomeruli, we measured STP in voltage clamp by presenting a series of optogenetic stimuli at different laser powers. Trials consisted of two stimulations separated by a delay. The first stimulation (stimulus 1) was 500 ms in duration with laser powers that varied between 0.1 and 1.7 mW in 0.1 mW steps. A second 250 ms stimulus was presented following a delay between 50 and 1,000 ms, with increased laser power corresponding to increased delays (stimulus 2). Likely due to a combination of membrane dynamics and optogenetic channel kinetics, we had less experimental control over the dynamics of the second stimulus. We empirically determined that using a power 0.25 mW larger than stimulus 1 produced the most reliable response with rates consistently between 50 and 80 Hz.

Data for the earliest collected KCs (n = 12) used a modified stimulation protocol. These experiments were performed before a laser realignment which altered PN spiking dynamics. Specifically, this protocol used a slightly higher range of laser power (0.4-2.25 mW) and a fixed delay of 250 ms between stimulus 1 and 2. Under these conditions, increased laser power produced a progressive prolongation of PN spiking past stimulus offset and introduced natural variability in the effective delay between stimuli. The realignment removed this prolongation and prompted the development of the protocol described above. Both protocols were designed and optimized to enable reliable STP parameter estimation while accounting for the dynamics of evoked PN spiking in each optical condition.

Trials were presented in a pseudo-randomized order. Each set consisted of one presentation of every power level, with the order of power levels randomized separately for each set. All connected glomeruli were stimulated at the selected power before proceeding to the next. We used an 8 s inter-trial interval and a minimum of 30 s between repeated stimulations of the same glomerulus. We aimed for 12 repeats per power level, but degradation of cell health or recording stability typically resulted in 4-8 repetitions. We interspersed current-clamp recordings after every fourth set of voltage-clamp trials. These recordings were not analyzed for STP dynamics but instead used the higher signal-to-noise ratio to monitor cell health and response latency.

Control experiments were performed on PNs to quantify spike rate dynamics and spatial resolution of photostimulation. Spike rate dynamics were measured using the same protocol described above for KC recordings. For lateral resolution, we stimulated a voxel whose edge was 1-2 μm from the border of the target glomerulus. For axial resolution, we presented stimuli above and below the focal voxel in 5 μm steps. We imaged Alexa Fluor 568 fills to determine glomerulus boundaries. PN recordings to quantify spike rate dynamics were performed regularly to validate consistency of the laser alignment and stimulation protocol across time.

### Cell type identification

We identified KC subtype using immunohistochemistry to visualize biocytin fills. Brains were fixed in 4% formaldehyde for 15 minutes, washed in 0.1 M PBS, and then blocked with 5% normal goat serum in 0.5% PBST. Brains were then incubated with mouse anti-Bruchpilot antibody (Developmental Studies Hybridoma Bank, nc82, 1:25 dilution) in blocking solution for 24 hours at 4°C on a nutator. Brains were washed again in PBST and incubated with goat anti-mouse Alexa Fluor 633 (1:250 dilution) and streptavidin Alexa Fluor 568 (1:500 dilution) for 24 hours at 4°C on a nutator. After washing in PBST, brains were mounted in Vectashield (Vector Laboratories) and imaged using a Zeiss LSM 880 confocal laser scanning microscope. KC subtype (γ, α’β’, or αβ) was determined based on axon anatomy.

KC subtype was inferred from physiology for six cells with unsuccessful biocytin fills. Inferences were made based on consistency across our recordings (including those not in the final dataset) and previously published results^50,75,76^. Two were identified as α’β’ due to high levels of baseline noise and a low spike threshold. αβ (n=1) and γ (n=3) were distinguished by differences in spike threshold and spike rate following current injection. Analyses based on KC subtype did not change when these cells were excluded.

### Network architecture

Networks were constructed with 50 PNs and 2,000 KCs that were modeled as conductance-based LIF neurons. Each KC received input from 6 randomly selected PNs. We did not model the modest connectivity biases reported in the connectome^32^. Varying the number of PN inputs per KC between 4 and 10 produced qualitatively similar results. KCs also received a combination of experimentally motivated feedforward and feedback inhibition^49,77^ that included subthreshold but no suprathreshold inhibition. All parameter values used for model simulations can be found in Table S1.

Because the membrane dynamics of an LIF neuron following spike generation are a key assumption that could influence temporal integration, we constructed three different model architectures in parallel. The primary model reported throughout the results was a multicompartment model where each PN synapsed onto an independent dendritic claw that was connected to a spike-generating soma by a conductance g_axial_. Feedforward inhibition was applied directly to each claw while feedback inhibition was applied to the soma. Membrane voltage was updated according to:

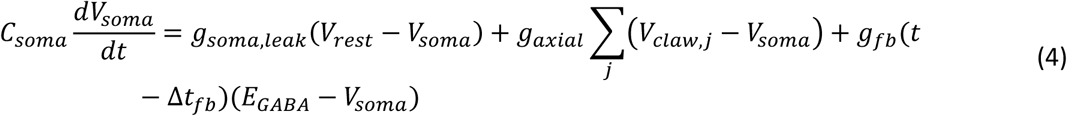

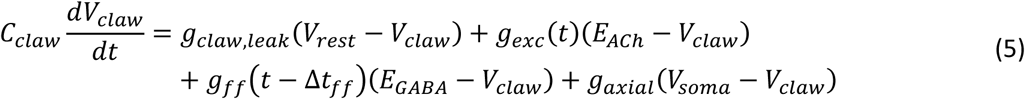

where C denotes the capacitance, V denotes the voltage, E denotes different reversal potentials, and g denotes different conductances. Inhibitory conductances were delayed Δt relative to the associated driving signal. Spikes were generated when V_soma_ > V_thresh_, triggering a reset of V_soma_ to V_reset_. Dendritic voltages were unchanged to allow for continued temporal integration. All parameter values are listed in Table S1.

Both of the alternative architectures were modeled as single-compartment models. In the first, a KC spike triggered a reset of its membrane voltage to V_reset_ for a refractory period t_refractory_. In the second, KC spikes instead triggered a transient increase in V_thresh_ that decayed back to baseline at the rate τ_thresh_. Membrane voltage for both models was updated according to:

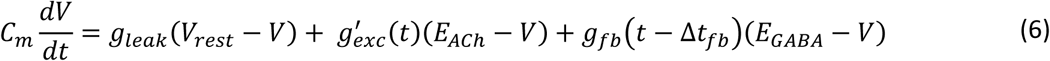

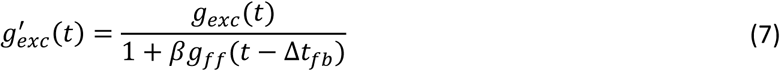

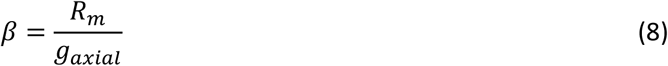

In these single-compartment models, feedforward inhibition was treated as a purely divisive, shunting current on each PN input with strength governed by β. All parameter values are listed in Table S1.

Initial parameter estimates for the model were obtained from published electrophysiology and analysis of data from this study. We estimated membrane resistance (R_m_) from 12 synapses in this study with the largest first EPSC amplitudes. We measured the average first EPSC amplitude from voltage-clamp recordings as well as the first EPSP amplitude from current-clamp recordings and computed R_m_ using Ohm’s law (R_m_ = 700 ± 100 MΩ). We estimated membrane capacitance (C_m_) using the hemibrain connectome. We extracted KC skeletons and computed the surface area assuming each segment was a perfect cylinder and assuming a specific capacitance of 2 μF/cm^2^.

Remaining parameters were initialized from published electrophysiological measurements (Table S1), and all parameters were adjusted within physiologically plausible ranges. Preliminary simulations indicated substantial parameter degeneracy, with changes in individual parameters compensated by others while preserving overall network activity and all qualitative conclusions. Reported results did not differ with parameter variability, conditional on overall E-I balance leading to 100-600 active KCs per odor.

### Synaptic dynamics

Model synapses were assigned experimentally constrained Tsodyks-Markram parameters (main text, equations 1-3). We implemented the model using its analytical solution which integrates synaptic state variables between presynaptic spikes to allow for continuous-time estimation of EPSC amplitudes. This implementation was used for both experimental fitting (see below) and network simulations. STP phenotypes were independently generated and assigned to each synapse using a Gaussian copula constructed from our experimental parameter estimates. First, we sampled a multivariate normal distribution with a correlation matrix estimated from our model fits to experimental data. These samples were transformed to uniform before mapping them to actual parameter values using the inverse cumulative distribution function of the experimentally measured marginal distributions.

To create networks with a reduced degree of STP heterogeneity, we created a pool of potential synaptic phenotypes using the same procedure. For within-KC heterogeneity networks, each KC randomly selected six phenotypes without replacement. These selected phenotypes were replaced in the pool for potential selection by the next KC. For within-KC homogeneity networks, each KC sampled a single phenotype which was assigned to all 6 synapses. Phenotypes were selected without replacement across KCs until the pool was depleted, at which point the pool was replenished to maintain maximal diversity across KCs.

We assigned the baseline EPSC amplitudes, A_0_, using a two-step procedure motivated by previously reported homeostatic regulation of KC excitability^78^. We empirically optimized this procedure such that the overlap in KC populations responding to two uncorrelated odors most closely matched the expected overlap from a binomial distribution. We first assigned A_0_ according to the exponential weighting function:

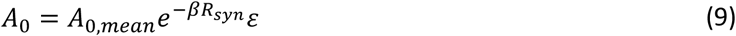

where A_0,mean_ = 4.5 pA, β = 0.6, R_syn_ is the steady-state response of the synapse to a 50 Hz spike train, and ε ∼ LogNormal(0, 0.1). For networks with no STP, the resulting A_0_ was used as the fixed-weight. After independently assigning A_0_ to each synapse, we secondarily scaled it to account for differences in the overall excitability of each KC. Each synapse was simulated with an identical 5 s, 50 Hz Poisson-distributed spike train. We then calculated the average EPSC across all spikes and synapses onto a given KC (R_KC_). We then multiplicatively scaled A_0_ for each synapse according to:

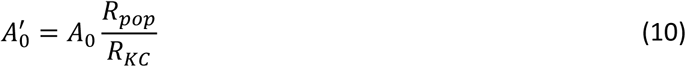

where R_pop_ is the average R_KC_ across the population. This normalization preserved relative differences in synaptic strength while reducing variability in overall excitability across KCs. Scaling factors were typically small (90^th^ percentile range: 0.91-1.09).

We used the assigned parameters of each synapse to convert PN spike trains into EPSC amplitudes. EPSC amplitudes were subsequently converted to point conductances by dividing each EPSC amplitude by the synaptic driving force (V_hold_-E_ACh_) where V_hold_ = −65 mV used experimentally and E_ACh_ = 0 mV. These point conductances were then convolved with a synaptic alpha function to generate continuous conductance waveforms. To determine the time constant of the alpha function, we extracted EPSC waveforms from the experimental recordings, normalized their amplitudes, and fit the average with an alpha function, yielding a time constant of 1.8 ms.

### Inhibition

Each network included experimentally motivated feedforward and feedback inhibition^49,77^. Feedforward inhibition was driven by PN input spikes and was specific to the associated excitatory conductance. Inhibitory conductances were delayed by 2 ms relative to excitation, had an alpha function time constant of 5 ms, and were scaled to 5% of the corresponding excitatory conductance. Feedforward inhibition included a simplified short-term facilitation, with a multiplicative gain of 1.5 and a time constant of 100 ms.

Feedback inhibition was driven by KC subthreshold voltage rather than population spiking, consistent with previous experimental observations^77^. We modeled this inhibition phenomenologically because the temporal dynamics of this inhibition are not fully characterized. Feedback inhibition was implemented as a band-pass filtered transformation of KC subthreshold voltage using a combination of a low-pass integration term (∼0.5 Hz) and a high-pass memory trace (∼3.5 Hz). Filtered activity was summed across the KC population with spatial weighting (space constant = 50 KCs) to approximate experimental observations that KCs most strongly inhibit themselves^74^. Gain was adjusted to match experimentally observed activity levels and applied to KCs as an inhibitory conductance after a delay of 10 ms. All attempts to incorporate feedback inhibition based on population spiking resulted in population dynamics similar to networks with no STP.

### PN odor responses

PN odor response PSTHs were parameterized to qualitatively reproduce experimentally measured data characterized by strong onset transients that decay to a more stable adapted rate^39,47,48^ (Table S1). Peak and adapted rates were sampled from exponential distributions with mean rates of 60 Hz and 40 Hz, respectively. These rates were clipped at maximum values of 250 Hz and 100 Hz, respectively. Onset latencies were sampled from a normal distribution with negative values clipped at 0 s. Time constants dictating the rate of adaptation were sampled from normal distributions for both the peak rate and a substantially slower decay of the adapted rate. PSTHs were generated using a difference-of-exponentials function with the latency to the peak rate defined as the onset latency plus 50 ms. All parameters were drawn with defined correlations relative to the sampled peak firing rate (Table S1).

Firing rates for step-function PNs were drawn from an exponential distribution with a mean rate of 50 Hz and included no variation in onset latency. Correlated odors were created by combining the parameters of a reference odor with an independent multivariate normal sample. The mixing coefficient determined the correlation strength while preserving the covariance structure of the response parameters.

Each network-odor pair was simulated for 20 independent trials. Although the primary KC population metrics stabilized with approximately eight trials, additional trials were included to support MBON learning analyses. Spike trains for each independent trial were generated by sampling ISIs from a gamma distribution. A shape parameter of κ = 5 was used unless otherwise specified based on analysis of unpublished data from our laboratory. Candidate ISIs were generated assuming the peak firing rate of each PN and a 2 ms refractory period.

Candidate spikes were then accepted using a standard thinning algorithm with probability 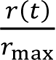, where r(t) is the instantaneous firing rate and r_max_ is the peak firing rate.

### MBON learning

A single MBON per network was modeled as a rate-based leaky integrator that received convergent input from every KC. Its dynamics were updated according to:

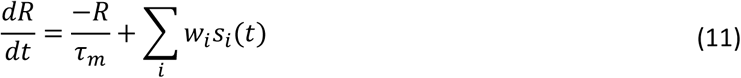

where R is the MBON activity, τ_m_ = 0.2 s is the membrane time constant, w_i_ is the weight between KC_i_ and the MBON, and s_i_(t) are the odor evoked spike times for KC_i_. All synapses were initialized with equal starting weights set to one.

Synaptic weights were updated after each training trial according to a learning kernel centered around the time of reinforcement. The kernel was averaged over all spikes fired by each KC with weights and a soft-bounded multiplicative weight update was computed proportional to that average:

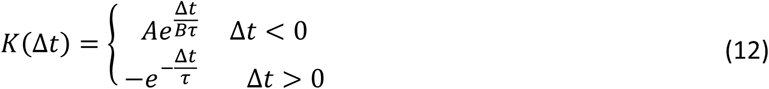

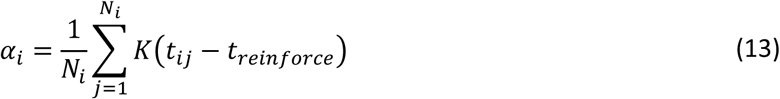

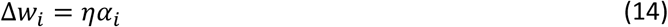

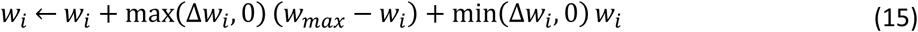

where K defines the learning kernel with parameters A =0.5, B = 2.5 and τ = 0.25 s that control the shape of the LTP arm relative to the LTD arm, N_i_ is the number of spikes fired by KC_i_, η = 25 is the learning rate, and w_max_ = 2w_0_ is the maximum weight allowed for a synapse. The kernel shape was empirically tuned using a parameter sweep to reproduce experimentally observed learning dynamics. Control and test MBON responses were computed using the same stimulus trials using either the initialized or trained weights, respectively.

## QUANTIFICATION AND STATISTICAL ANALYSIS

### PN spike detection and characterization

PN spikes recorded during control experiments were detected using an automated algorithm followed by manual verification. Individual traces were band-pass filtered between 100 and 1,500 Hz, and spikes were detected using an amplitude threshold on the derivative of the trace selected for each cell. Detected spikes were visually inspected and manually corrected.

Axial resolution was measured relative to the boundary of the target glomerulus. Boundaries were estimated based on Alexa Fluor 568 fills imaged under 2p illumination. In instances where cells did not fill, we conservatively estimated boundaries based on visual inspection of mVenus expression in the antennal lobe. Experiments were discarded if glomerulus boundaries could not be confidently identified. Because responses did not differ for stimulation locations above versus below the glomerulus, measurements from both directions were combined when binning distances in 5 μm increments.

We evaluated variability in the evoked ISI distribution using a time-rescaling analysis. Instantaneous firing rates were estimated by smoothing spike trains with a Gaussian kernel (σ = 20 ms) and each ISI was transformed according to the time-rescaling theorem:

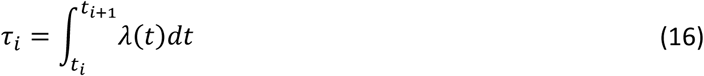

where λ(t) is the instantaneous firing rate. Time-rescaled ISIs were compared with spike trains generated from measured firing-rate trajectory using Poisson-distributed ISIs.

### EPSC detection

EPSCs were identified and measured for each voltage-clamp trial using custom-written MATLAB scripts. Initial event detection was performed using an automated algorithm based on the derivative of the filtered trace. Event detection was subsequently refined using extensive manual curation to remove false positives and identify missed events. Because presynaptic spike times were inferred from KC responses rather than recorded directly from PNs, manual event identification was guided by the statistical properties of PN spiking characterized from control recordings. Trials where EPSCs could not be confidently detected were discarded. Because event identification depended on evaluating response consistency across repeated trials within a connection, manual EPSC detection was performed using the full set of trials rather than blinded individual traces. Only connections with at least 25 analyzed trials were included, with only 3 reported synapses having less than 30 trials and 12 synapses with less than 40 trials.

We first identified the EPSC evoked by the initial spike in each trial, as the initial response amplitude strongly constrains the characterization of synaptic dynamics. In PN recordings, onset latencies were stereotyped across trials (standard deviation < 15 ms after accounting for inverse correlation between rate and latency), providing a reliable estimate of first EPSC time when comparing across KC voltage-clamp trials. Mapping trials and interspersed current-clamp trials provided a second source of latency information, as the higher signal-to-noise ratio more clearly identified response onset compared to voltage-clamp trials. Because exceptions were observed in PN recordings, this information was used conservatively rather than enforced as a strict criterion.

EPSC identification across the full stimulus used the statistical regularity of evoked PN spike trains to constrain the expected timing and occurrence of synaptic responses. EPSC detection was jointly evaluated based on the voltage-clamp trace and the inferred sequence of ISIs relative to the expectation of highly stereotyped spiking established in PN recordings (ISI SD < 10 ms). Importantly, response detection was not conditional on observing a successful neurotransmitter release because release failures (EPSC = 0) provide meaningful information characterizing the response of a KC to a PN spike. Thus, putative failures and other small-amplitude EPSCs were considered a response when their inclusion improved consistency with the ongoing sequence of inferred ISIs. Conversely, when multiple candidate responses occurred within an implausibly short interval relative to surrounding ISIs, the response assignment that produced the most consistent ISI sequence was retained. Because PN regularity was not perfect, with variability partially locked to the pulsed stimulation protocol, these criteria guiding spike time inference were applied conservatively. Smoothing over 3-5 successive ISIs removed some of this variability and was considered as a secondary source of information for resolving ambiguous cases. Importantly, early spikes are the most influential and informative for estimating STP dynamics, so criteria were applied most conservatively over the first 5-10 EPSCs, and trials were discarded if these ambiguities could not be confidently resolved.

The same statistical analyses used to characterize PN spike train dynamics were applied to inferred spike trains from KC recordings. These analyses provided a quantitative comparison of spike timing statistics between directly recorded PN responses and inferred presynaptic activity during voltage-clamp experiments.

### Fitting Tsodyks-Markram model to data

We fit the Tsodyks-Markram model of STP to measured EPSC data using a published algorithm that uses a Bayesian inference framework to estimate the marginal posterior distributions using Markov chain Monte Carlo sampling^44^. Briefly, the algorithm compares observed synaptic responses with predictions from sampled parameters while incorporating response uncertainty into the likelihood function. For each connection, the algorithm was provided with inferred EPSC times, measured EPSC amplitudes, and estimates of response variability. For fitting, we used the default priors and MCMC settings provided in the published MATLAB implementation.

To obtain an estimate of EPSC variability, we clustered trials based on the inferred onset spike rate. Bin edges were manually selected to include approximately 10–15 trials per bin. Initial bin boundaries were preferentially placed at discontinuities in the rate distribution, and additional bins were introduced as needed to achieve the desired sample size per bin. Synaptic responses were averaged across trials to estimate the mean and standard error. Because trials exhibited small differences in firing rate, each EPSC was associated with uncertainty in both amplitude and spike timing. The Bayesian fitting algorithm explicitly accounts for uncertainty in response amplitude but not spike timing. We therefore assumed that the small temporal uncertainty contributed to the measured amplitude variability, an approximation that is consistent with the locally linear dynamics assumed by the Tsodyks–Markram model. To obtain the spike times associated with each averaged EPSC, we first converted ISIs for each trial into an instantaneous rate. Representative spike times for each bin were reconstructed from the average instantaneous rate trajectory. Once the train of EPSCs had fewer than five trials contributing to the average, the response was terminated and further EPSCs discarded.

We implemented the fitting algorithm several times per connection using different preprocessing steps to evaluate estimation robustness. First, onset rate measurements varied depending on how many EPSCs the instantaneous rate was smoothed over (default = 5, range = 3-8). Second, we varied how many EPSCs were included in the model input for stimulus 1 (default = 16, range = 8-25). This range balanced allowing synaptic dynamics to approach steady state before the delay separating stimulus 1 and stimulus 2 while excluding the final portion of stimulus 1 where accumulated differences in spike rate resulted in increasing response uncertainty that could lead to overfitting as well as increased uncertainty in EPSC inference. Third, we varied how many EPSCs were included for stimulus 2 (default = 1, range = 1-5). This second stimulus primarily functioned to constrain estimates of τ_fac_, with the first EPSC amplitude being the most informative. Spike rates during stimulus 2 were less tightly controlled experimentally, increasing uncertainty and disproportionately weighting parameter estimation toward the 60–80 Hz range typical of the second stimulus.

We defined the peak of the marginal posterior distributions as the parameter values describing each connection. Preliminary analyses and initial parameter recovery simulations showed that this value was substantially more reliable across data preprocessing and initial conditions compared to the joint MAP estimate or a more standard least-squares approach. These approaches necessarily find the best-fitting parameters, but improvements were typically less than 1% (R^2^ and RMSE) compared to reported fits using the marginal posterior modes. For most connections, predicted dynamics were robust across preprocessing iterations and the default preprocessing scheme was kept. For a subset of connections, fits were broadly convergent but produced minor differences in predicted dynamics. In these cases, preference was given to solutions that most accurately reproduced the stimulus 2 response. Although all responses contribute equally to the likelihood function, this EPSC provides the strongest constraint on facilitation parameters, and its influence can otherwise be diluted by the larger number of observations from stimulus 1. Connections that failed to produce reliable parameter estimates were discarded.

Fit quality was measured as the RMSE between the measured and predicted EPSC amplitudes. A_0_ was fixed at the estimated amplitude when measuring the RMSE of the reported fits. For the no STP comparison, we identified the fixed-weight amplitude that best fit the data using a least-squares approach. To compare against alternative parameter sets, we generated 100,000 sets of randomly generated parameters. Amplitude was a free parameter that was scaled to best fit the recorded data using least-squares fitting. This amplitude fitting included a bias term, λ = 3, that was used to slightly overweight the first EPSC. We also generated model parameters using the Gaussian copula to match the distribution of the population fits and found no differences from fully random sampling.

### Analysis of synaptic organization

We performed hierarchical clustering on the estimated Tsodyks-Markram fit parameters. All parameters except U_0_ were first log-transformed before clustering, reflecting their log-normal distributions and allowing clustering to capture relative rather than absolute differences in parameter values. We then z-scored each parameter to account for differences in parameter magnitude that would lead to τ_fac_ and A_0_ dominating the measured distances. Clustering used the cosine distance and an average linkage function. Compared to alternatives, this combination of transformations and clustering produced both qualitatively similar functional clusters and the highest silhouette scores (s = 0.56), except for untransformed variables (s = 0.78) where clustering was dominated by the high magnitudes of τ_fac_ and A_0_.

Synaptic organization was quantified using the intra-class correlation using a one-way ANOVA approach for partitioning the variance based on KC identity, KC subtype, and PN identity. ICC was measured as:

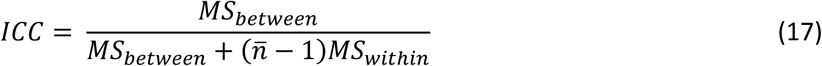

where MS_between_ is the between-KC mean square, calculated as variance of the KC means weighted by number of synapses. MS_within_ is the within-KC mean square, calculated as the residual variance around KC means. ñ is the average number of connections within each group. The null distribution was generated using 10,000 shuffles of cell identity while maintaining the same distribution of synapses per KC. Confidence intervals were measured using 10,000 bootstraps. For the multivariate ICC, PCA was performed on the log-transformed and z-scored parameter set.

### Analysis of model KCs

Model KCs were considered active if they averaged at least 2.5 spikes per stimulus (> 0.25 Hz). KC PSTHs were constructed by binning spike times in 1 ms increments and convolving with a 250 ms Gaussian kernel. Peak latencies were estimated as the time where the PSTH reached its maximum rate. For the PSTH of *in vivo* KC odor responses, we excluded the first 1 s of the response for peak latency estimates to exclude a strong onset transient, following the original published analysis^37^.

Temporal coverage was measured using a normalized participation ratio which, quantifies the dispersion of KC peak latencies across the duration of the stimulus and is defined as:

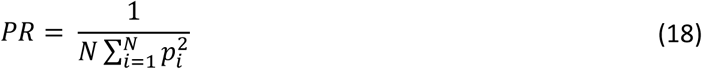

where p_i_ is the fraction of KCs with a peak latency in bin i and N is the total number of bins. Bin size was fixed at 500 ms and results were robust to variations in bin width between 200 and 1,000 ms.

Response sparseness was calculated for single KCs using the Treves-Rolls sparseness measure. Spike times were binned in 250 ms bins and sparseness was calculated as:

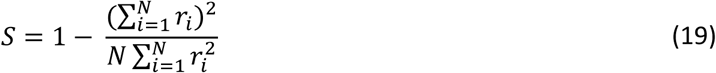

where r_i_ is the rate in bin i and N is the total number of time bins. While magnitude did vary with bin size, both networks and analysis of *in vivo* KC recordings had similar shifts.

Response reliability was measured using a cross-validation of estimated peak latencies. The 20 trials per network-odor pair were separated into groups of 10 trials and the peak latency was estimated for each. Reliability was then defined as Spearman’s rank correlation between the peak latencies measured for each set.

We decoded time after stimulus onset using a Bayesian likelihood decoder previously developed and described in detail for analysis of *in vivo* KC odor responses^37^. Briefly, spike trains were binned into 500 ms time windows and the expected spike counts per bin were estimated for each KC using leave-one-out cross-validation. For each observation, the likelihood of the observed spike counts was computed for each time bin assuming an independent Poisson model. Log likelihoods were summed across the population to generate a posterior probability distribution for time predictions. Decoding accuracy was quantified as the RMSE between the posterior mean estimate of decoded time and the true time, averaged across time bins.

To determine whether KCs had consistent peak latencies across odors, temporal preference was quantified as the variance of the peak latency across odor responses for each KC. Because estimating variance requires multiple odor responses per cell, these sparsely active networks were simulated with 150 odors (17.4 ± 2.4 responses per cell). Significance was assessed using a permutation test where peak latency values were randomly reassigned across KCs and odors, preserving the latency distribution while eliminating any cell-specific temporal preference. Empirical p-values were computed from 1,000 permutations using a lower-tailed test for reduced variance compared to the null distribution. Multiple comparisons were corrected using the Holm-Bonferroni method.

Unless otherwise specified, all modeling results are reported for 20 independent network replicates that were each simulated with the same set of 50 odors. For each replicate, metrics were first averaged across odors, and reported values represent the mean ± standard deviation across the 20 network replicates. Averaging across networks first resulted in small increases in standard deviations but does not affect interpretation of any results.

### Model PN temporal information

We first evaluated temporal decoding of model PN responses using the same Bayesian decoder described for the KC population. The decoder was applied to the simulated PN spike trains used as input to the network, with decoding performed across the population of 50 PNs.

For a more precise measure of temporal information contained within PNs, we calculated the Fisher information of the population. PSTHs were down sampled to 500 ms to match the temporal resolution of the Bayesian decoder, although results did not qualitatively change with different bin sizes. A lower bound on temporal uncertainty was then calculated according to:

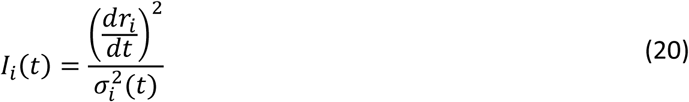

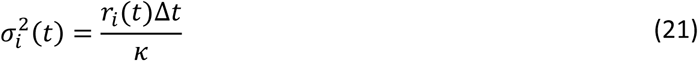

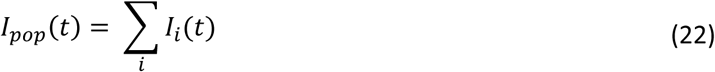

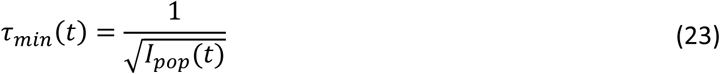

where r_i_(t) is the PSTH of PN_i_, σ_i_^2^(t) is the expected variance of the spike count within a bin, κ is the shape parameter governing ISIs drawn from a gamma distribution, and τ_min_ is the Cramér-Rao bound that represents the lower bound on temporal estimation implied by the PN population.

### Pattern separation analysis

Pattern separation was measured using 5 network replicates with each network simulated with 10 independently generated odor families at each correlation value. Temporal information was discarded and the population response was converted to a binary activation vector indicating whether each KC responded to a given odor. Pattern separation was quantified by measuring the population overlap using the Dice-Sørensen coefficient:

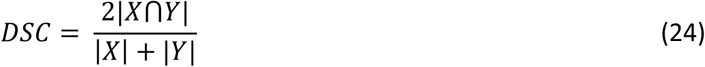

where X and Y are the sets of active KCs for two odors. For visualization, histograms include all pairwise odor comparisons across all odor families and networks. For comparing STP and no STP models, we first averaged across networks for each odor family resulting in 10 independent odor family replicates per correlation level. Note that this averaging differs from other analyses that instead treated networks as the independent replicate. Similar means and variances were observed when networks rather than odors were treated as independent replicates.

---

**Figure S1.**
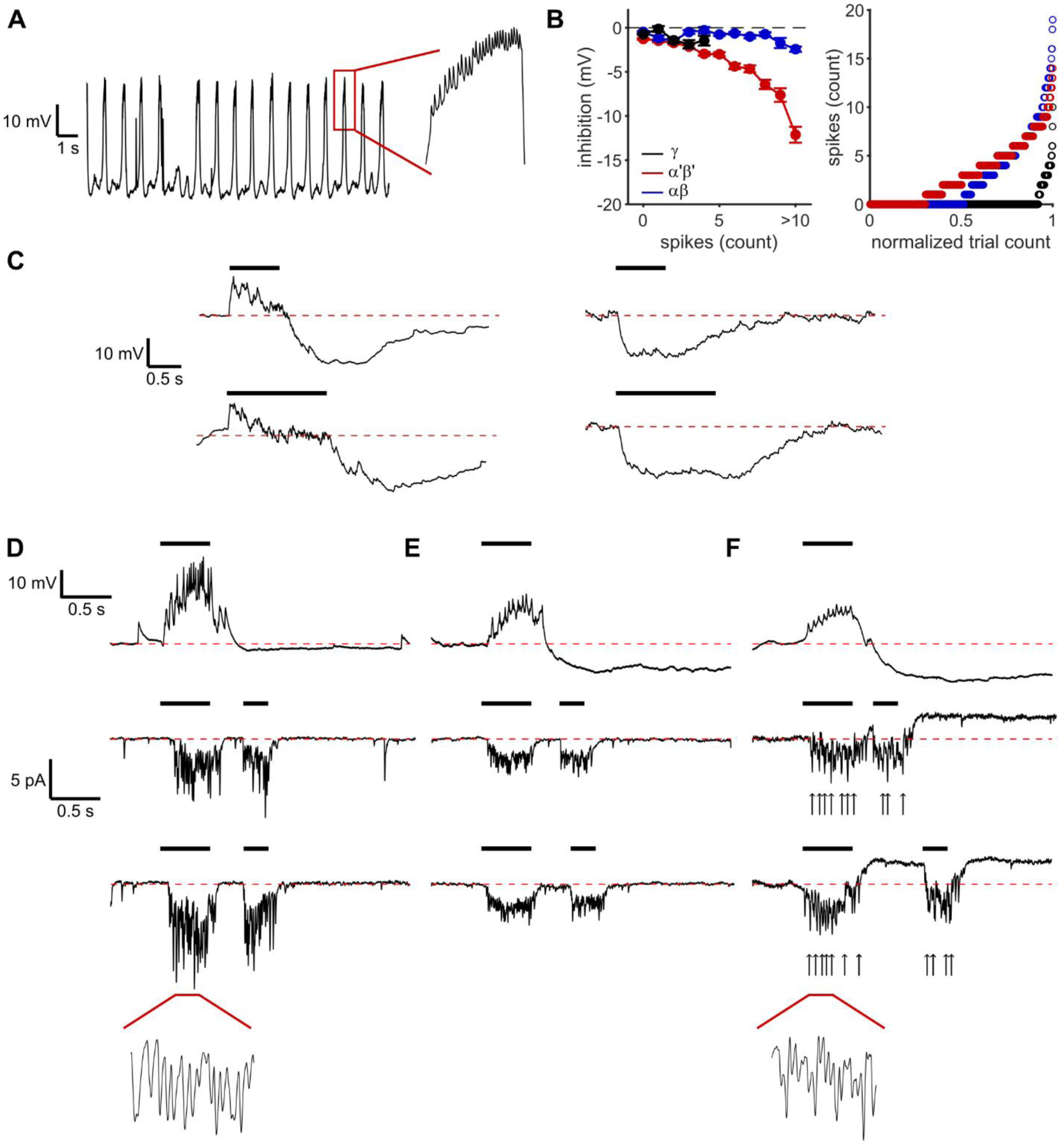
Inhibition is minimal during single-glomerulus stimulation in voltage clamp. **(A)** PN recording in the presence of picrotoxin (2.5 μM) and the absence of stimulation shows the emergence of population-wide oscillatory activity that impeded STP measurements. **(B)** Quantification of inhibition (left) and spike count (right) measured in KCs during 2-photon stimulation of single glomeruli at the highest laser power levels (>1.5 mW). Inhibition was measured as the magnitude of hyperpolarization in the 1 s following stimulus offset. Example trials are shown in panel D. The small post-stimulus hyperpolarizations in γ and αβ KCs are known to be non-GABAergic^50^. **(C)** Recordings during direct activation of the mushroom body calyx. GH146-Gal4 expresses in PNs and APL, so these traces represent combined activation of PN axon terminals and APL-mediated inhibition. All tested KCs (n = 6) had detectable inhibition. The γ KC (left) shows a combination of PN excitation and APL inhibition. The αβ KC on the right was a rare instance where no connections with CsChrimson-expressing PNs were detected. **(D)** Example traces from an αβ KC during single-glomerulus stimulation at a high laser power (1.8 mW). Spikes were common in current clamp (top) but were suppressed during voltage-clamp trials at the same laser power (middle and bottom). The inset zooms in on a 250 ms portion of the stimulus 1 response. Contrasting with the panel F inset shows that unclamped spikes and high-amplitude EPSCs are distinguishable. EPSCs typically have a slower and more symmetric trajectory. **(E)** Example traces from an α‘β‘ KC during single-glomerulus stimulation at a high laser power. High spike rates and prominent inhibition were observed in current clamp (top, 1.6 mW), both of which are suppressed during voltage clamp (middle, 1.6 mW). **(F)** Example traces from an α‘β‘ KC with space-clamp failures during voltage clamp, shown as positive evidence that spiking and inhibition are detectable in our preparation. High spike rates and strong inhibition are present during current clamp (top, 1.5 mW). Both unclamped spikes and prominent inhibition are detectable during voltage clamp for this KC. Unclamped spikes had a characteristic shape consisting of a sharp deflection in the current trace followed by a large overshoot, as seen in the inset. Unclamped spikes occurred almost exclusively in α‘β‘ KCs and were excluded from analysis.

**Figure S2.**
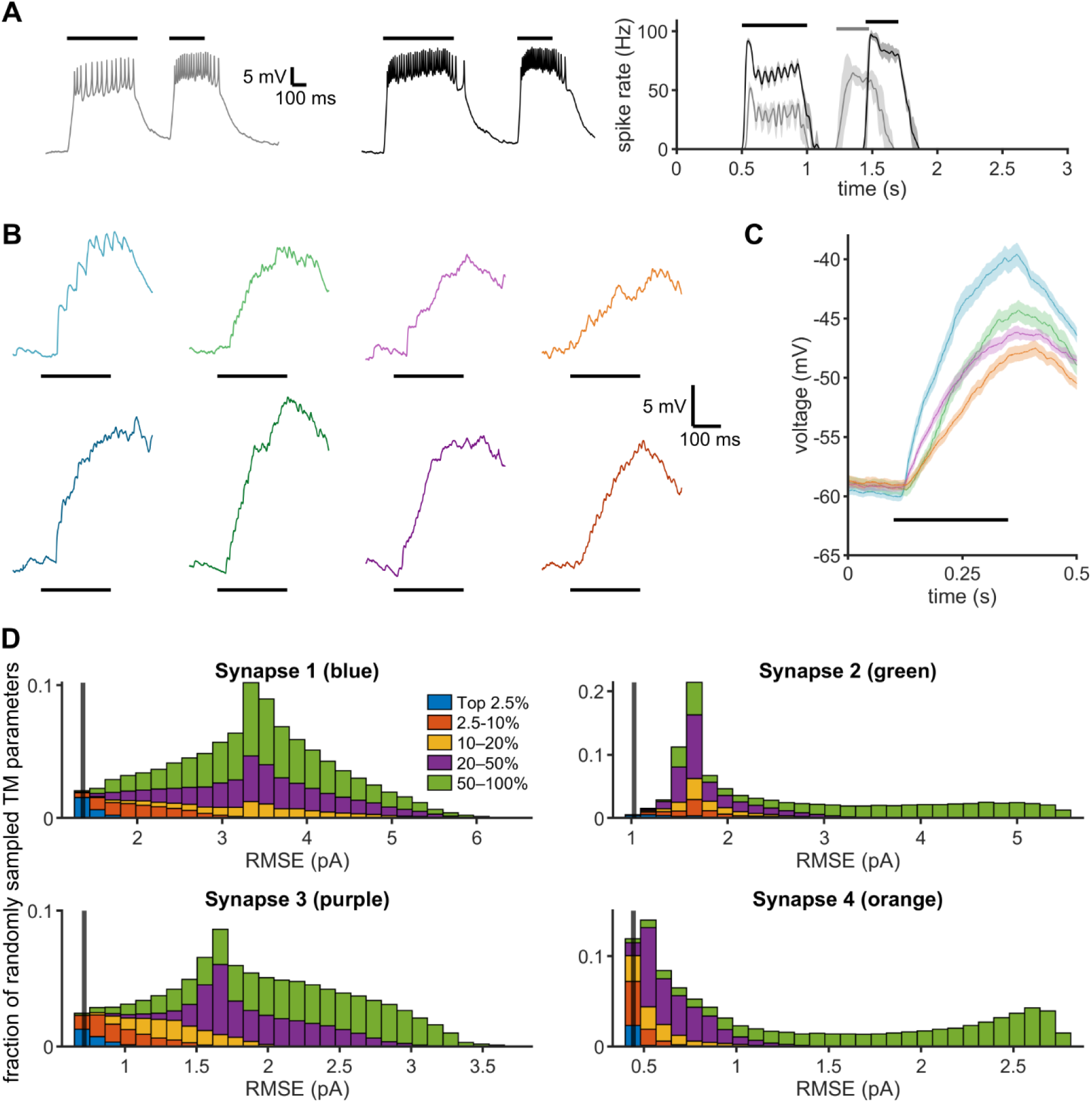
Example KC mapping data and fit quality. **(A)** Our stimulation protocol was updated mid-project to account for changes in PN spiking statistics caused by a substantial realignment of the laser. Specifically, the original protocol had a fixed delay between stimulus 1 and stimulus 2. PN spiking consistently extended into the delay period with a duration correlated with spike rate, which created a natural (but not explicitly controlled) distribution of inter-stimulus intervals. The realignment provided experimental control over stimulus offset, and the updated protocol introduced variable delays, with example PN responses depicted on the left and the average PSTH across trials (n = 10 trials, mean ± SEM) on the right. Both stimulation protocols were optimized within experimental limits to enable accurate estimation of Tsodyks-Markram parameters. **(B)** Example current-clamp responses to mapping stimuli for the γ KC shown in main text Figures 1 and 2. Although laser power was fixed across mapping trials, voxel locations affected PN spike rate, with panels showing responses to low (top) and high (bottom) PN spike rates. **(C)** Average (mean ± SEM, n = 22 blue, 18 green, 23 purple, and 24 orange) voltage response to mapping trials for the glomeruli shown in panel B. Note that inter-stimulus interval and spike rate could vary substantially across mapping trials and therefore were used strictly to assess qualitative agreement between current-clamp and voltage-clamp data. **(D)** Histograms show the RMSE between data and model fits for 100,000 randomly selected combinations of Tsodyks-Markram parameters. Each panel corresponds to a synapse in panel A and the main text Figures 1 and 2, indicated in the panel title. Black line shows the RMSE of the true estimated fit. Bars are colored based on the average marginal posterior density and binned into percentiles.

**Figure S3.**
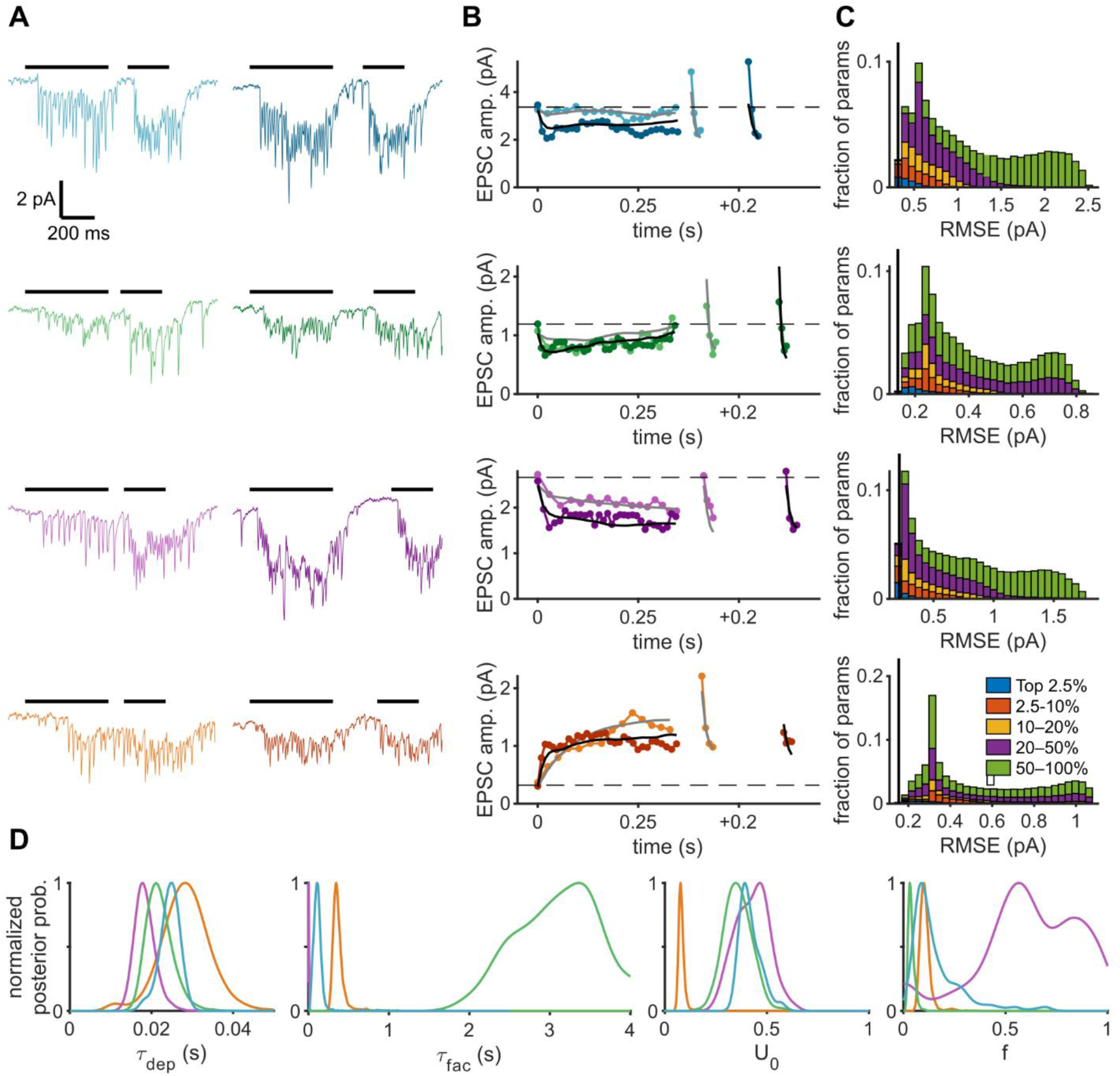
Example STP dynamics from an α’β’ KC. **(A)** Example voltage-clamp trials from an α’β’ KC at two different laser powers using the updated protocol. Each row shows a different PN synapsing onto the same KC. **(B)** Average EPSC amplitudes and Tsodyks-Markram predictions for two different spike rates for the same KC. The stimulus 1 response is truncated at 0.3 s and stimulus 2 is plotted relative to the stimulus 1 offset. Error bars are not shown to improve visual clarity. **(C)** Comparison of the RMSE between predicted and measured EPSCs. Vertical black lines indicate the RMSE for the estimated parameters. Histograms represent 100,000 randomly selected combinations of Tsodyks-Markram parameters. Colors represent the average posterior density of each parameter set and binned by percentiles. **(D)** Posterior probability distributions for each connection shown in Panel C.

**Figure S4.**
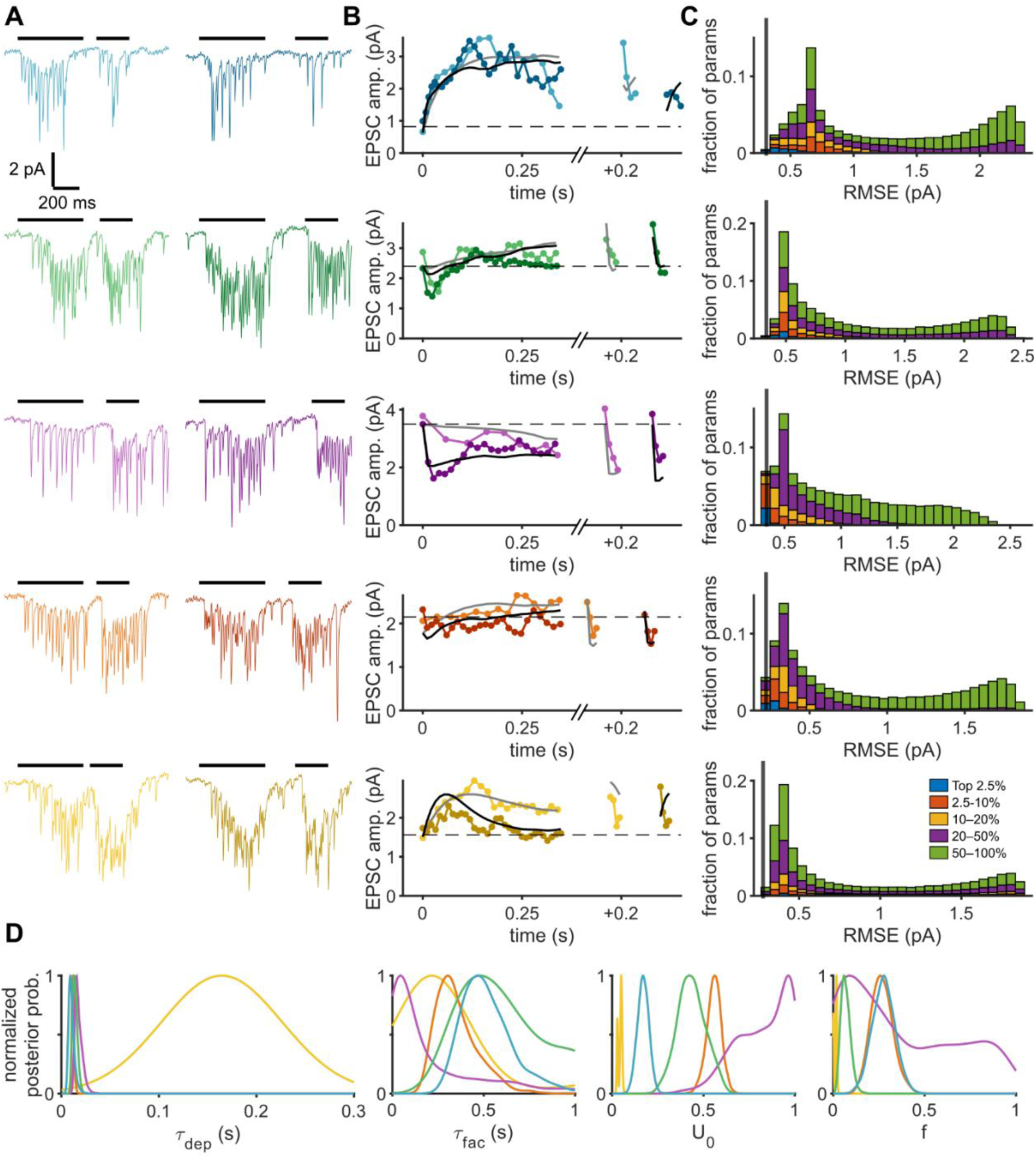
Example STP dynamics from an αβ KC. **(A)** Example voltage-clamp trials from 5 PNs synapsing onto the same αβ KC at two different laser powers. **(B)** Average EPSC amplitudes and model fits for the connections in Panel A at two different spike rates. The stimulus 1 response is truncated at 0.3 s, and stimulus 2 is plotted relative to the stimulus 1 offset. Error bars are not shown for visual clarity. Note that this KC has two representative examples of commonly observed complex STP phenotypes. Green has a depressing-facilitating phenotype, which consistently had stronger onset depression than the model could capture while nonetheless following the overall envelope. Yellow shows a facilitating-depressing synapse that the model could typically reproduce more accurately. **(C)** Comparison of RMSE between measured EPSCs and model predictions. Vertical black lines indicate the RMSE from the estimated parameters. The histogram shows RMSEs across 100,000 randomly selected combinations of parameters. Bars are colored by the average posterior density of each parameter set and binned by percentiles. **(D)** Posterior probability distributions for each synapse shown in Panel C.

**Figure S5.**
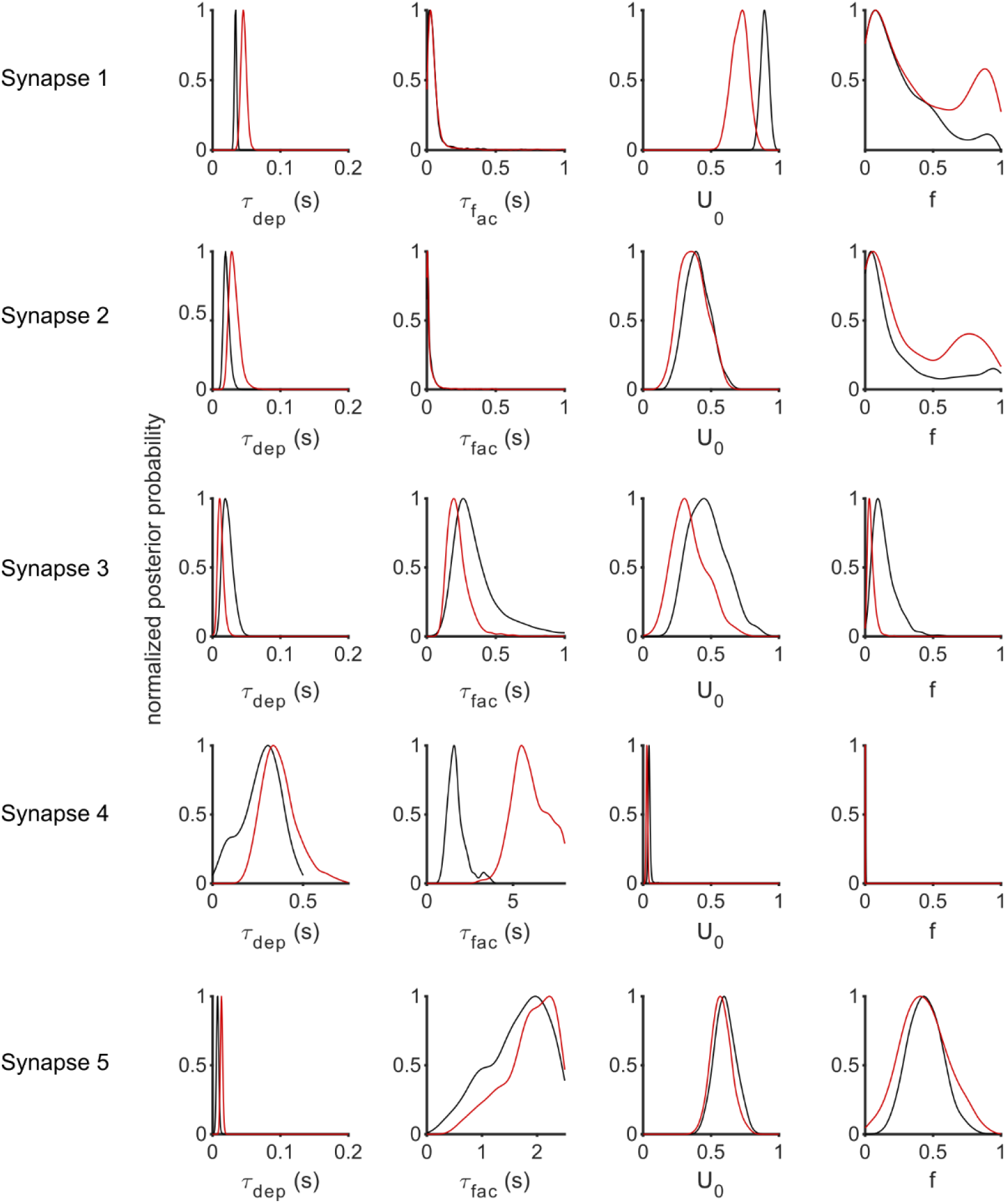
Tsodyks-Markram estimates are reliable across independent fits. In five experiments, two locations within large and spatially extended glomeruli were selected for STP characterization, with data analyzed independently for each site and blind to the matching glomerulus identities. Posterior distributions are shown for all parameter estimates. The fit shown in black was used for all analyses due to a more central glomerulus location that led to a greater number of analyzed trials and a larger range of inferred PN spike rates. Note that the divergent fit of τ_fac_ for synapse 4 is partially due to magnitude (2 s vs. 5 s), both of which indicate a long-lasting facilitation. This discrepancy likely reflects limitations in stimulation of the red location, where stimulus 2 failed to evoke spikes in most trials, reducing the resolution of this estimate.

**Figure S6.**
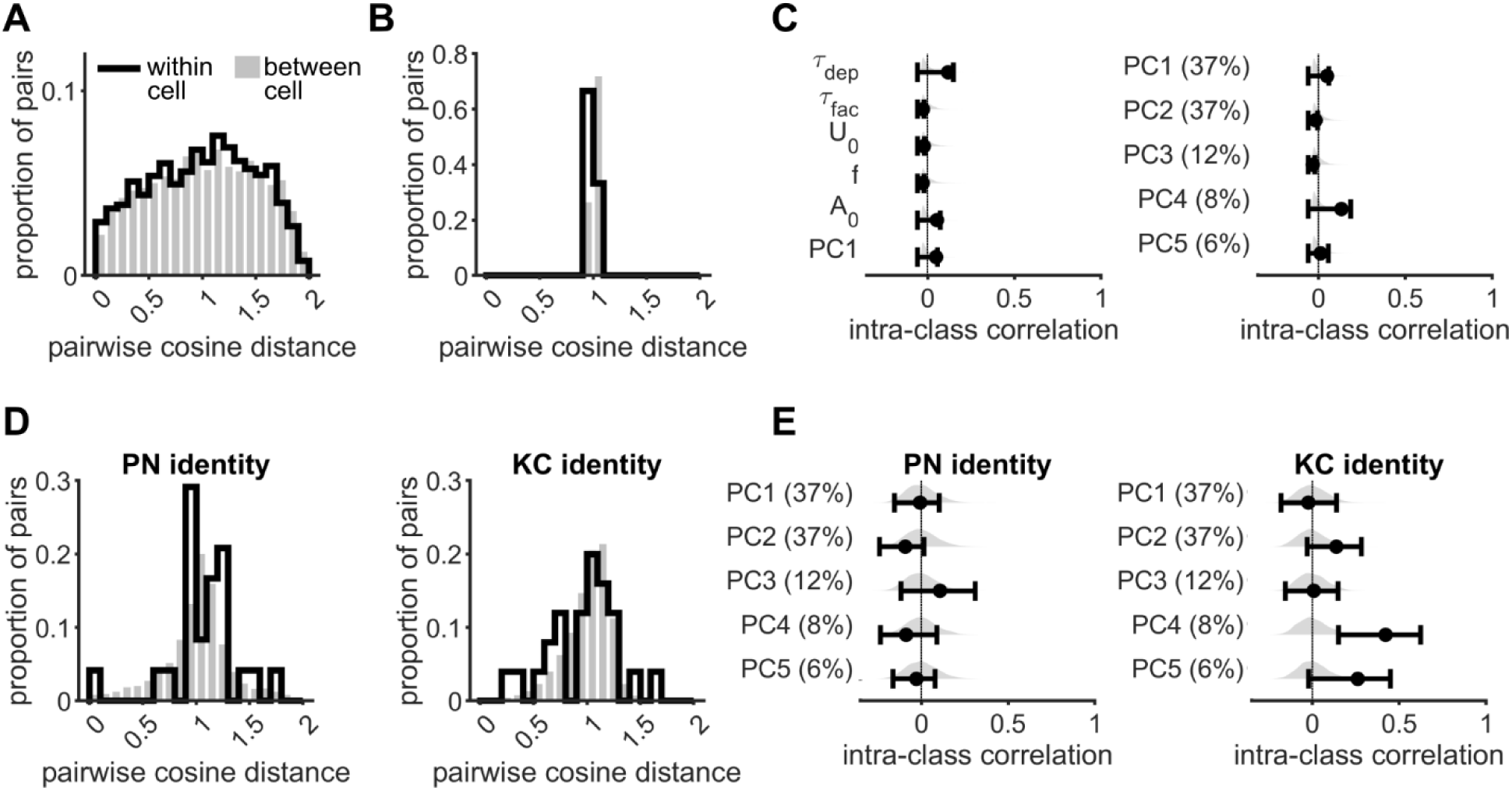
KC subtype cannot explain STP organization. **(A)** Distribution of pairwise distances between all recorded synapses, separated into pairs that are part of the same KC subtype (γ, α‘β‘, αβ) or different subtypes (p = 0.11, permutation test with 10,000 shuffles). **(B)** Pairwise distances were averaged across all pairs sharing the same KC subtype. Between-subtype differences (gray shaded bars) were generated by shuffling KC subtype identity and recomputing (p = 0.65, permutation test with 10,000 shuffles). **(C)** ICCs for data grouped by KC subtype, showing results for individual Tsodyks-Markram parameters (left) and each principal component (right), with percentages indicating the percent of variance explained for each PC. **(D)** Pairwise distances were averaged across all within-glomerulus (left) or within-KC (right) pairs. Between-PN (p = 0.16) or KC distances (p = 0.84) were generated by shuffling synapse identity (permutation tests with 10,000 shuffles). **(E)** ICCs for all principal components based on PN identity (left) or KC identity (right). Percentages indicate the percent of variance explained for each PC.

**Figure S7.**
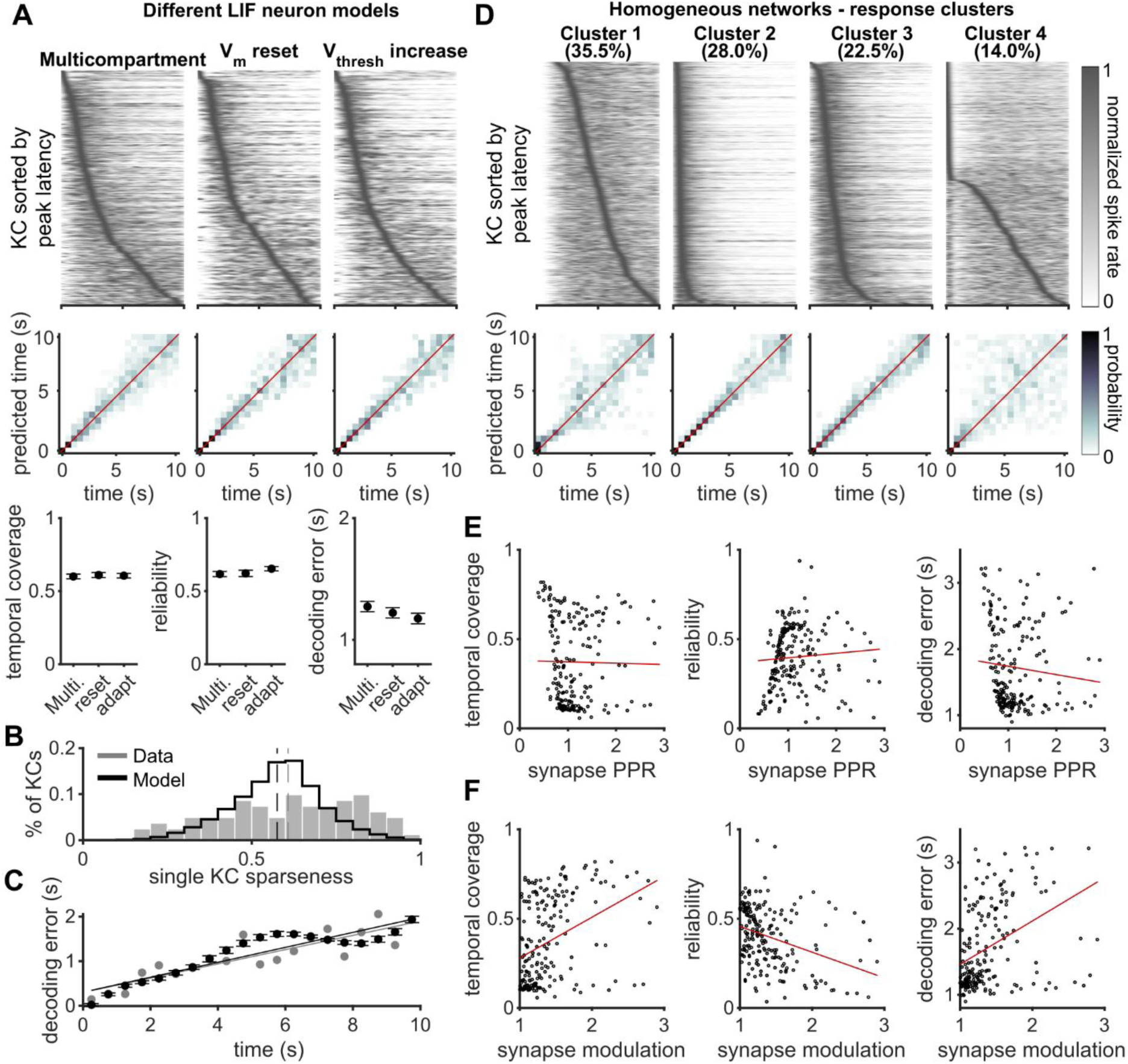
Additional quantification of network temporal dynamics. **(A)** Networks with different assumptions about how the voltage of an LIF neuron responds to a spike were constructed in parallel to test the potential effects of this nonlinearity on temporal integration and coding. Plotted are example KC responses with the same STP organization and same simulated odor using these different neuron models and quantified across replicates (n = 20). **(B)** Temporal sparseness was measured for individual KCs in both published *in vivo* odor responses and networks simulated here (p = 0.50)^37^. **(C)** Decoding error plotted as a function of stimulus time for previously published data and simulated networks (mean ± SD, n = 20). Fits were not significantly different (p = 0.28, Kolmogorov-Smirnov test). Experimental data were reproduced from Kropf et al. (2026)^37^, used under a CC BY 4.0 license. **(D)** Networks with homogeneous STP (n = 200) were manually classified based on visual inspection of population PSTH dynamics. **(E)** Metrics quantifying temporal structure compared to the 50 Hz paired-pulse ratio of each network with homogeneous STP (n = 200 networks, p > 0.05 for all three panels). **(F)** Same as (E) but x-axis instead shows synapse modulation, where facilitating synapses (i.e. PPR > 1) are plotted the same as panel E but depressing synapses are instead plotted as the inverse of the PPR. All correlations were significant (temporal coverage: r = 0.38, p < 0.001, reliability: r = −0.34, p < 0.001, decoding error: r = 0.40, p < 0.001).

**Figure S8.**
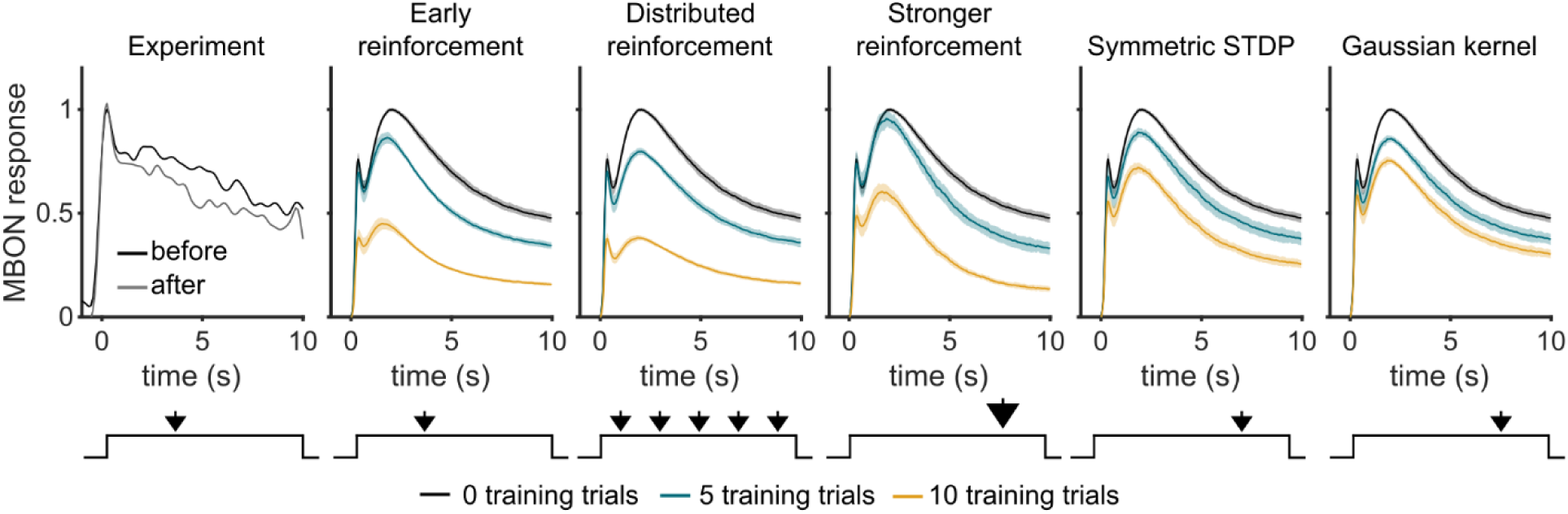
MBON learning with alternative reinforcement schemes. MBON learning was assayed using different reinforcement schemes. In each panel, black is untrained response, blue is after 5 learning trials, and yellow is the response after 10 learning trials. Experimental data showing MBON response with reinforcement at 3.5 s was reproduced from Kropf et al. (2026)^37^, used under a CC BY 4.0 license.

**Table S1:** Network model parameters for three different LIF neuron implementations.

| **Parameter** | **Multicompartment** | **Voltage reset** | **Dynamic threshold** |
|---|---|---|---|
| *KC properties* |  |  |  |
| E<sub>ACh</sub> | 0 | 0 | 0 |
| E<sub>GABA</sub> | -70 mV | -70 mV | -70 mV |
| V<sub>rest</sub> | -55 mV | -55 mV | -55 mV |
| V<sub>threshold</sub> | -45 mV | -45 mV | -45 mV |
| V<sub>reset</sub> | -60 mV | -55 mV | - |
| ΔV<sub>thresh</sub> | - | - | 15 mV |
| τ<sub>ΔVthresh</sub> | - | - | 10 ms |
| t<sub>refractory</sub> | 2.5 ms | 2.5 ms | 2.5 ms |
| R<sub>m,soma</sub> | 1.5 GΩ | 600 MΩ | 600 MΩ |
| R<sub>m,claw</sub> | 5 GΩ | - | - |
| C<sub>m,soma</sub> | 14 pF | 28 pF | 28 pF |
| C<sub>m,claw</sub> | 2 pF | - | - |
| g<sub>e</sub> gain | e<sup>1.275</sup> | e<sup>1.1</sup> | e<sup>1.1</sup> |
| g<sub>e</sub> alpha function time constant | 1.8 ms | 1.8 ms | 1.8 ms |
| g<sub>axial</sub> | 3 nS | 3 nS | 3 nS |
| *g<sub>i,ff</sub> (feedforward inhibition)* |  |  |  |
| Gain relative to excitation | 0.05 | 0.8 | 0.6 |
| Delay relative to excitation | 2 ms | 2 ms | 2 ms |
| STP gain per spike | 1.5 | 1.7 | 1.5 |
| STP decay time | 100 ms | 100 ms | 100 ms |
| Alpha function time constant | 5 ms | - | - |
| *g<sub>i,fb</sub> (feedback inhibition)* |  |  |  |
| Space constant | 50 KCs | 50 KCs | 50 KCs |
| Gain relative to V<sub>m</sub> | 0.1 | 1 | 1 |
| Delay | 10 ms | 10 ms | 10 ms |
| Integration time constant | 50 ms | 50 ms | 50 ms |
| Memory time constant | 750 ms | 750 ms | 750 ms |
| Low-pass coefficient | 0.75 | 0.5 | 0.65 |
| High-pass coefficient | 2.5 | 1 | 1 |
| *PN odor responses* |  |  |  |
| κ | 5 | 5 | 5 |
| Peak rate (mean) | 60 | 60 | 60 |
| Adapted rate (mean) | 40 | 40 | 40 |
| Adapted rate correlation to peak | 0.4 | 0.4 | 0.4 |
| Onset latency (mean) | 0.1 | 0.1 | 0.1 |
| Onset latency (SD) | 0.1 | 0.1 | 0.1 |
| Latency correlation to peak rate | -0.5 | -0.5 | -0.5 |
| Peak adaptation rate (mean) | 2.5 s | 2.5 s | 2.5 s |
| Peak adaptation rate (SD) | 0.5 s | 0.5 s | 0.5 s |
| Stable adaptation rate (mean) | 100 s | 100 s | 100 s |
| Stable adaptation rate (SD) | 15 s | 15 s | 15 s |

